# Collateral fitness effects of mutations

**DOI:** 10.1101/820068

**Authors:** Jacob D. Mehlhoff, Frank W. Stearns, Dahlia Rohm, Buheng Wang, Erh-Yeh Tsou, Nisita Dutta, Meng-Hsuan Hsiao, Courtney E. Gonzalez, Alan F. Rubin, Marc Ostermeier

## Abstract

The distribution of fitness effects (DFE) of mutation plays a central role in constraining protein evolution. The underlying mechanisms by which mutations lead to fitness effects are typically attributed to changes in protein specific activity or abundance. Here, we reveal the importance of a mutation’s collateral fitness effects, which we define as effects that do not derive from changes in the protein’s ability to perform its physiological function. We comprehensively measured the collateral fitness effects of missense mutations in the *E. coli TEM-1* β-lactamase antibiotic resistance gene using growth competition experiments in the *absence* of antibiotic. At least 42% of missense mutations in *TEM-1* were deleterious, indicating that for some proteins, collateral fitness effects occur as frequently as effects on protein activity and abundance. Deleterious mutations caused improper post-translational processing, incorrect disulfide-bond formation, protein aggregation, changes in gene expression, and pleiotropic effects on cell phenotype. Deleterious collateral fitness effects occurred more frequently in *TEM-1* than deleterious effects on antibiotic resistance in environments with low concentrations of the antibiotic. The surprising prevalence of deleterious collateral fitness effects suggests they may play a role in constraining protein evolution, particularly for highly-expressed proteins, for proteins under intermittent selection for their physiological function, and for proteins whose contribution to fitness is buffered against mutations with deleterious effects on protein activity and protein abundance.

**Significance Statement:** Mutations provide the source of genetic variability upon which evolution acts. Deleterious protein mutations are commonly thought of in terms of how they compromise the protein’s ability to perform its physiological function. However, mutations might also be deleterious if they cause negative effects on one of the countless other cellular processes. The frequency and magnitude of such collateral fitness effects is unknown. Our systematic study of mutations in a bacterial protein finds widespread collateral fitness effects that were associated with protein aggregation, improper protein processing, incomplete protein transport across membranes, incorrect disulfide-bond formation, induction of stress-response pathways, and unexpected changes in cell properties. Our results suggest that deleterious collateral fitness effects may be an important constraint on protein evolution.

## Main Text

Fitness is one of the most important concepts in evolutionary biology. Despite its importance, fitness can be difficult to measure because it is a combination of many traits that interact in complex ways (1). Accordingly, researchers generally estimate fitness using fitness components traits like fecundity and survivorship that are thought to be the result of the most important traits affecting fitness. Although these estimates have been very useful for evolutionary research, they tell us little about the underlying mechanisms of these fitness components. To bridge this gap, the “functional synthesis” approach (2) combines molecular genetics mechanisms with fitness estimates from evolution and ecology to understand the molecular details of fitness components. We suggest that such research could also benefit from an “afunctional synthesis” a consideration of the innumerable ways that a protein must remain afunctional in order to maintain organismal fitness. For example, an enzyme catalyzing an intermediate step in glycolysis is afunctional with respect to being an inhibitor of DNA polymerase; mutations that cause such inhibition would assuredly be deleterious. While acquisition of the inhibitory function in this example may seem extraordinarily unlikely, this low probability is countered by the innumerable ways that proteins must remain afunctional to avoid deleterious fitness effects within the complex milieu of the cell.

Protein misfolding and misinteractions are likely sources for such deleterious fitness effects. Misinteractions can originate from a protein either in its native state (3-6) or a misfolded conformation (7-9). For example, sets of missense mutations that cause misfolding of the yellow fluorescent protein (YFP) cause up to a 3.2% decrease in yeast growth rate in a concentration-dependent manner (9). A decrease in fitness as a result of toxic, insoluble protein is an example of what we are defining here as *collateral fitness effects*. Collateral fitness effects are those effects that do not derive from changes in the ability of the gene to perform its physiological function. In the case of YFP, the mutational effects on fitness derive from the insoluble YFP and not from the corresponding loss of soluble, functional YFP because jellyfish YFP has no physiological function in yeast. Such collateral fitness effects are in contrast to *primary fitness effects* – the focus of the functional synthesis – which arise from changes in the ability of the gene to perform its physiological function (e.g. through changes in protein specific activity or active protein abundance). Mutations with deleterious collateral fitness effects cause the protein to do something it “should not” do, with adverse consequences for fitness. These two components of the fitness effect of mutation are not mutually exclusive. A mutation’s fitness effects may be the result of only primary fitness effects, only collateral fitness effects, or a combination of the two. The sign of primary and collateral fitness effects can be opposite, which sets up the possibility of fitness trade-offs. For example, a mutation can have beneficial primary effects yet remain deleterious for fitness due to larger, deleterious collateral fitness effects.

Delineating fitness effects into these two categories is useful because primary and collateral fitness effects manifest in different ways. Mutations have primary fitness effects only when fitness is impacted by changes in the protein’s ability to perform its physiological function. Primary fitness effects do not exist unless the protein is under selection for its physiological function. They also do not exist in regimes where organismal fitness is buffered from mutational effects on protein activity/abundance due to an excess of protein activity, the action of chaperones, or other mechanisms (10-13). For example, most enzymes are well above the threshold in activity that the cell requires, so small, deleterious effects on protein cellular activity might have no effect on fitness. In contrast, collateral fitness effects can exist in both these situations because they do not require changes in the ability of the protein to perform its physiological function or even that the protein’s physiological function has fitness relevance. Another difference lies in how deleterious primary and collateral fitness effects can be compensated by secondary mutations. For example, deleterious primary fitness effects that decrease specific activity can be compensated by an increase in protein abundance. In contrast, deleterious collateral fitness effects that arise from misfolding/misinteraction can be alleviated by a decrease in protein abundance.

We were interested in how much collateral fitness effects contribute to the distribution of fitness effects (DFE) of mutation. The DFE (in particular the distribution of deleterious mutations) plays a central role in constraining protein evolution, yet the functional basis behind the DFE continues to be debated (14). One approach to studying the DFE is to leverage large-scale, systematic measurements of mutational effects in experiments called Deep Mutational Scanning (DMS) (15, 16). In these experiments, large sets of mutant alleles are subject to selective pressure. The effects of the mutations are quantified by deep sequencing the alleles before and after selection. Many DMS studies use phenotypes or protein properties as a proxy for fitness (often outside the protein’s native context) with limited relevance for organismal fitness landscapes and protein evolution (17-20). DMS studies using growth competition as the selective pressure have measured organismal fitness effects (e.g. (13, 18, 21-24)) but have not delineated collateral fitness effects. Fitness effects were interpreted as arising solely from mutational effects on protein specific activity/abundance and how these physiochemical traits map onto the level of organismal fitness. The frequency and magnitude of collateral fitness effects is essentially unknown but is central to understanding how misfolding, misinteractions, and potentially other effects shape the DFE and protein evolution. Here, we use DMS to measure the frequency and magnitude of collateral fitness effects in the antibiotic resistance protein TEM-1 β-lactamase by conducting growth competition experiments in the absence of antibiotic (i.e. in the absence of primary fitness effects).

## Results

### Deleterious collateral fitness effects of mutations are common in TEM-1 β-lactamase

The key to measuring collateral fitness effects is to perform experiments under conditions where the protein’s physiological activity is irrelevant to organismal fitness. We chose to examine the fitness effects of mutations in the antibiotic resistance gene *TEM-1* β-lactamase in the absence of β-lactam antibiotics. TEM-1 with its N-terminal signal sequence (preTEM-1) is exported to the periplasm of *E. coli*, whereupon the signal sequence is proteolytically removed to yield the mature protein. TEM-1 provides β-lactam antibiotic resistance by hydrolyzing the β-lactam ring of antibiotics such as ampicillin (Amp), which interfere with cell wall synthesis. TEM-1 has no other known physiological role, so the antibiotic’s absence makes TEM-1 superfluous. Thus, we assume that fitness effects of mutations observed in a growth competition experiment in the absence of β-lactam antibiotics are collateral fitness effects. Although antibiotic resistance genes, which are beneficial in the presence of antibiotics, can have fitness costs in the absence of the antibiotic, such fitness trade-offs are not the subject of this experiment. We asked whether mutations cause a change in fitness relative to cells containing the wildtype allele when *both* are evaluated in the absence of the antibiotic. We reason that such effects will likely occur in the presence of the antibiotic as well and are an important (but hidden) component of the fitness effects of mutations.

We desired a TEM-1 expression system that allowed inducible, moderately high expression of TEM-1 without appreciable fitness effects or significant changes to the *E. coli* transcriptome. Plasmid-borne *TEM-1* is natively expressed from a strong, constitutive promoter in *E. coli* (e.g. as found on plasmid pBR322) and is one of the most abundant proteins in the periplasm (**Fig S1A**,**B**). We placed *TEM-*1 under the IPTG-inducible *tac* promoter on a plasmid with a *p15A* origin (pSkunk1), which has a lower copy number than pBR322. We utilized NEB 5-alpha cells that overexpress LacI to strongly repress the *tac* promoter in the absence of IPTG. We chose this experimental design to guard against losing deleterious alleles during library creation and propagation that preceded the growth competition experiment. The use of pSkunk1 and 1 mM IPTG provided about 2.7-fold higher levels of active TEM-1 than pBR322 (**Fig. S1C**). At most, this level of expression would be expected to have a fitness cost of 0.06% due to allocation of ribosomes to TEM-1 production (25). Any fitness effect of induction of TEM-1 expression was smaller than could be detected in our growth assay (**Fig. S2**). Western blotting detected the mature protein but did not detect preTEM-1 (**Fig S1B**), suggesting that our level of TEM-1 expression did not overburden protein export or precursor processing. TEM-1 expression caused almost no changes to the cell’s transcriptome (**Fig. S1D**). Thus, TEM-1 expression was moderately high to best provide the opportunity to observe collateral fitness effects, but not too high such that TEM-1 expression itself caused significant fitness or phenotypic effects.

We constructed libraries designed to contain all possible single-codon substitutions in *TEM-1*. We subjected these libraries to a growth competition experiment in the absence of β-lactam antibiotics. At the start of the growth competition experiment, we diluted uninduced, exponentially-growing cultures of the libraries into LB media containing IPTG. Cultures were allowed to grow exponentially for ∼10 generations at 37°C. We determined the frequencies of the wildtype and mutant alleles at the beginning and end of the experiment by deep sequencing using Enrich2 (26). From these frequencies, we determined the fitness values, which correspond to the mean growth rate of cells containing a mutant allele relative to the mean growth rate of cells containing the wildtype allele. We combined the results for synonymous codons because most fitness measurements among synonymous codons were statistically equivalent. Growth assays on a few of the rare instances in which one allele seemed to produce a dramatically different fitness effect than its synonyms showed that such instances were artifactual (**Figure S3**). From the fitness (*w*), we calculated the collateral fitness effects of each mutation as the selection coefficient, *s* = *w* − 1.0.

We found that collateral fitness effects for *TEM-1* are surprisingly frequent and can be large in magnitude. We measured the fitness effects of 94.9% (5428/5720) of the possible nonsynonymous mutations in TEM-1. Replica experiments indicated that most fitness values significantly different from 1.0 were reproducible (**Figure S4).** While many fitness values were near-neutral, 21.6% (1113/5157) of missense mutations caused fitness effects in both replica experiments (*P*<0.01, *Z*-test) (**Fig 1A**). A total of 18.1% (932/5157) of missense mutations caused fitness effects at *P*<0.001 in both replica experiments, and all of these were deleterious. We were able to detect reproducible, deleterious effects on growth rate as small as about 1% (*s* ranged from −0.011 to −0.84). Missense mutations 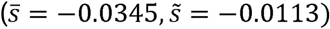 were more deleterious than nonsense mutation 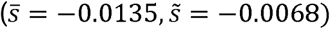 and synonymous mutations 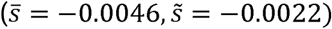. By assuming that fitness measurements for neutral mutations will be symmetrically distributed about a selection coefficient of 0.0 and that no mutations have a beneficial effect, we estimate that 42.0% of missense mutations in TEM-1 cause more than a 1% decrease in fitness (**Fig. 1B**). This is a lower bound on the frequency of deleterious mutations. Mutations with deleterious fitness effects |s| > 4 × 10^−8^, which is calculated from the inverse of the effective population size of *E. coli*, will constrain TEM-1 evolution (28). Such mutational effects are ∼10^5^ times smaller than we can measure in our DMS experiment. Our DMS technique does not allow us to distinguish very small deleterious fitness effects from truly neutral mutations and thus cannot fully capture the mutational constraints on TEM-1’s evolution.

**Figure 1.**
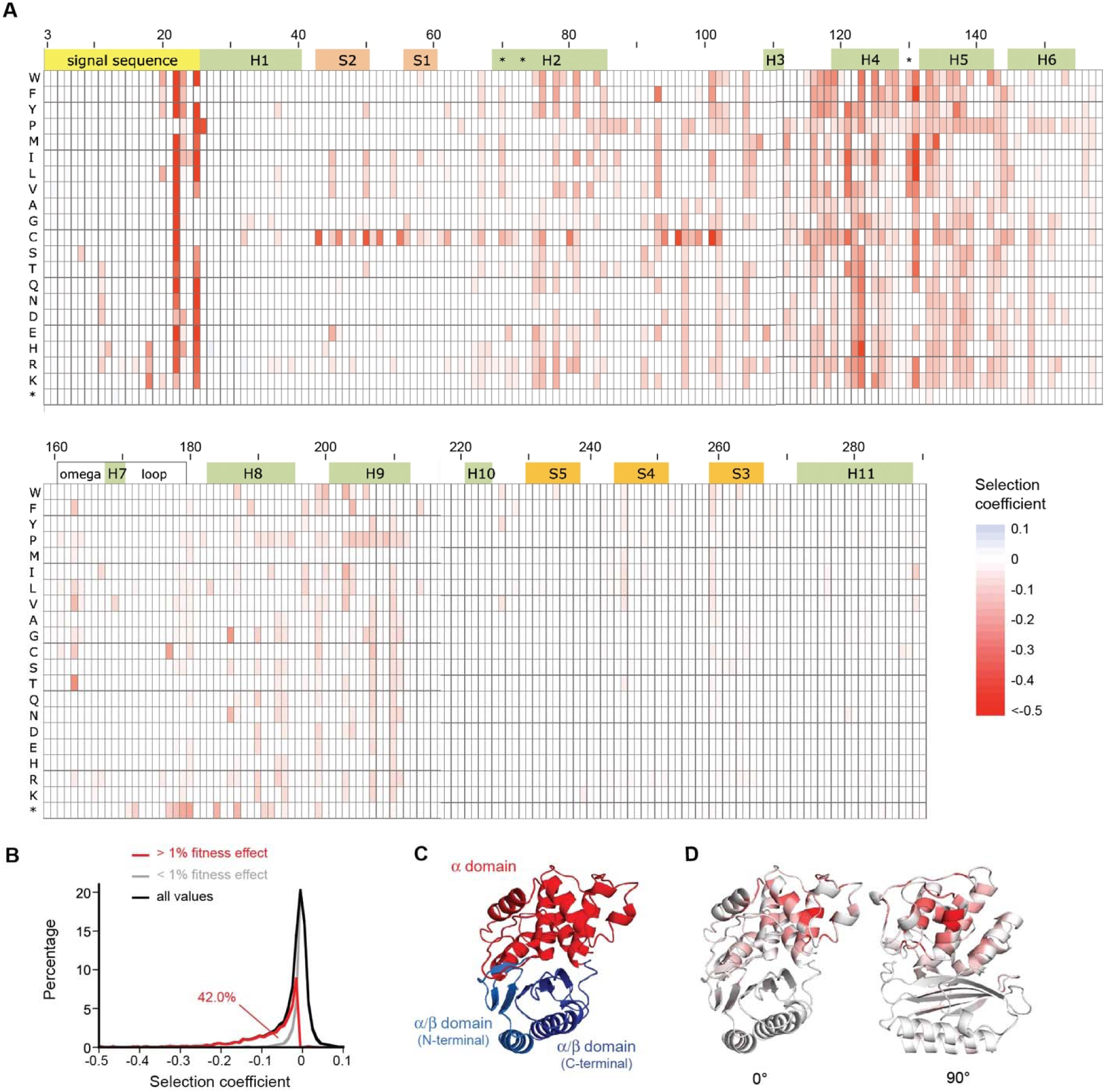
Collateral fitness effects of mutations in TEM-1 β-lactamase. (**A**) Heat map of weighted mean selection coefficients for mutations that caused fitness effects (*P*<0.01 in both replica experiments). These selection coefficients reflect the mean growth rate over ten generations post-TEM-1 induction relative to cells lacking a TEM-1 mutation. The Ambler numbering system (27) for class A β-lactamases is used. Regions corresponding to the signal sequence (yellow), α-helices (green), β-strands (orange), the Ω-loop (white) and three key active site residues (*) are shown. A heat map showing all fitness values is provided as **Fig. S5** and tabulated fitness values are provided as **Data S1**. (**B**) Distribution of fitness effects (DFE) for missense mutations indicating the estimated portion of missense mutations that cause a greater than 1% decrease in fitness in the absence of Amp. Structure of TEM-1 (PDB ID 1XPB) (29) showing (**C**) the α domain and discontinuous α/β domain and (**D**) the median fitness values for mutations at each position (white, no effect; red, increasing magnitude of deleterious effect).

TEM-1’s α domain (residues 69-212) was conspicuously prone to deleterious collateral fitness effects compared to the α/β domain (residues 26-68 and 213-290) (**Fig. 1C,D**). The frequency of deleterious missense mutations in the α domain (29.0%) was 5.5-fold higher that of the α/β domain (5.2%) for fitness values *P*<0.001 in both replicas. The median fitness effect in the α domain (−0.0269) was 4.6-fold larger than that in the α/β domain (−0.0058) (*P*<0.001; Mann-Whitney). α-helices 4 and 5 and the end of the signal sequence were particularly prone to large deleterious fitness effects. Mutations on the protein surface (defined as residues with >20% solvent accessible surface area), were less prone to deleterious effects than buried residues (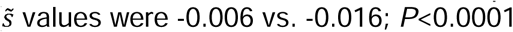; Mann-Whitney; mean values were −0.015 and −0.039). This result is consistent with the observation that surface residues evolve faster than residues buried in the core of the protein (30). Mutation to cysteine caused the highest mean and median deleterious fitness effects, followed by mutations to aromatic residues (**Fig. S6A**). Amino acid size was highly predictive of the mean fitness effect (*r*^2^ = 0.83), provided mutations to glycine, proline, and cysteine were excluded (**Fig. S6B**,**C**). Twenty-two nonsense mutations (7.7%) were deleterious in both replica experiments (*P*<0.01), with half of these (including the nine most deleterious) within the region between residues 170 and 195 (**Figs.1A** and **S7A**). Nonsense mutations at the beginning of the gene were not beneficial, which confirms that TEM-1 expression does not have a detectable fitness cost in our experiment.

### Monoculture and co-culture growth experiments confirm the collateral fitness effects

We constructed a set of 37 mutants to confirm their collateral fitness effects and investigate their mechanisms. These included twelve mutants in the signal sequence, eight mutants to or at cysteine residues, eight mutants in α-helices of the α domain, and nine nonsense mutations in the region 170-195. This set included S70N, K73C, S124G, and several nonsense mutations that lacked a fitness effect. S70 and K73 are key active site residues at which missense mutations inactivate the enzyme.

We monitored the growth of monocultures expressing the TEM-1 mutants over 10 generations of exponential growth as measured by the cultures’ optical density (OD). We used the starting and final OD values to calculate a mean growth rate (relative to cells lacking a mutation) over ten generations post-induction, analogous to how we calculated fitness effects in the DMS experiments. We also measured the steady-state growth rate over the timespan of 4-6 hours after induction. These growth assays supported the deleterious effects for most mutations (**Fig 2, Fig S7B**). They also revealed that deleterious fitness effects did not fully manifest until 2-3 hours after the induction of TEM-1 expression (**Fig S8**). This delay is consistent with the expectation that transcription, translation, and accumulation of TEM-1 take time to reach steady state in the cell. Thus, the deleterious collateral fitness effects measured by DMS and the mean growth rate measured by monoculture growth are underestimates because these fitness measurements are solely based on the allele frequency or culture density, respectively, at the beginning and end of the 10-generation growth period.

**Figure 2.**
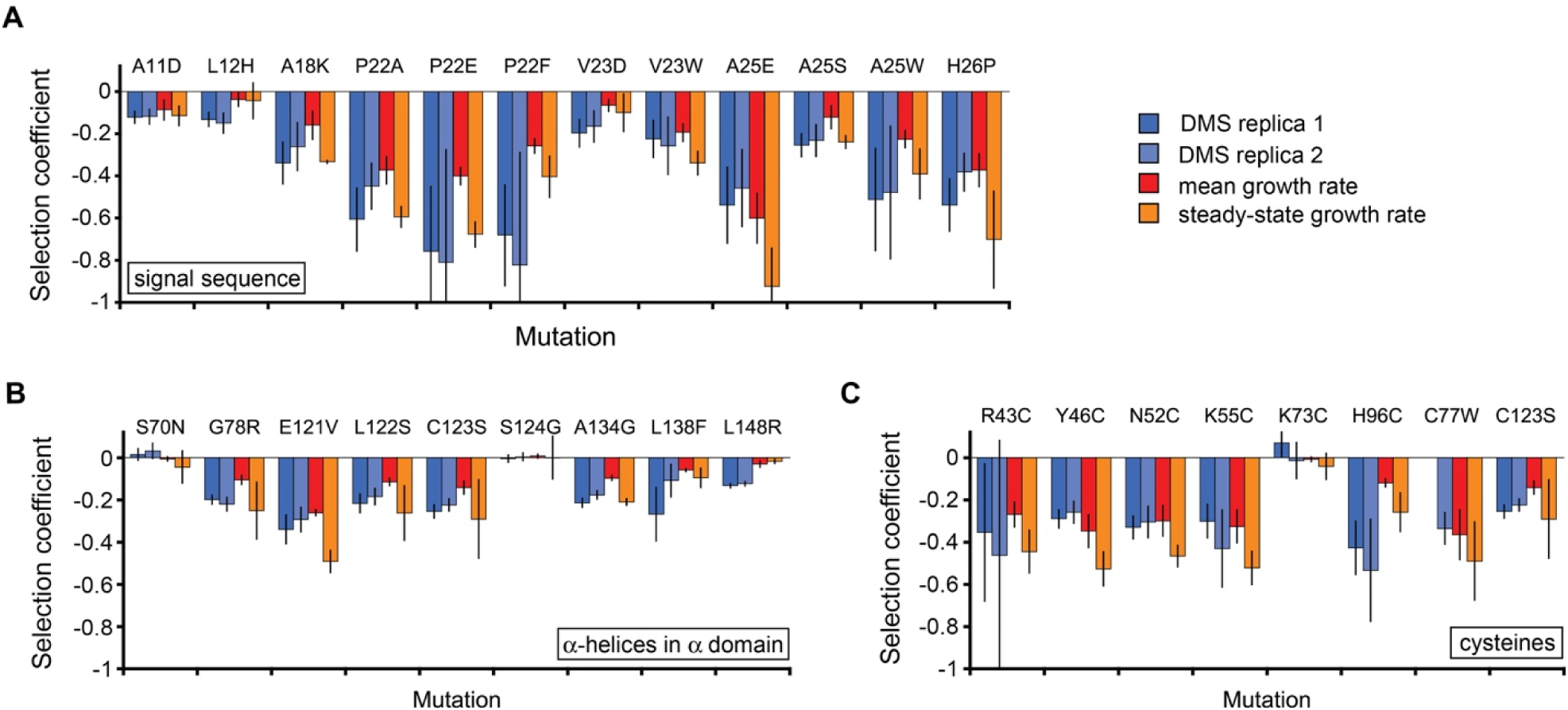
Confirmation of collateral fitness effects. Selection coefficients for cells with TEM-1 mutations (**A**) in the signal sequence, (**B**) in α-helices 2, 4, 5 and 6 of the α domain, and (**C**) involving cysteine. Selection coefficients were measured by deep mutational scanning (blue, two biological replicates), mean growth rate over 6 hours post-induction (red; *n*≥3), and steady-state growth rate 4-6 hours post-induction (*n*≥3). Error bars represent 99% confidence intervals. For the mean and steady-state growth rates, cells expressing the wildtype and mutant proteins were grown in separate flasks for ten generations of wildtype growth as monitored by OD-600nm. Representative growth data provided as **Fig. S8**. DMS fitness values for S124G exclude data for the GGT codon for Gly (see **Fig. S3**).

We found that the deleterious fitness effects of mutations as measured in the growth assay were sometimes less than those measured by DMS. For two such mutants (A134G and L148F), we measured the fitness effects of the mutations in co-culture by two different methods. These results were more consistent with the fitness measurements by DMS (**Fig S9**). We do not know the origin of the differences between the DMS and growth assay measurements. A plausible explanation for the difference is that the relationship between culture OD and cell concentration is affected by the mutation (i.e. differences would occur if the mutations affect the cells ability to scatter light). Mutations might affect cell division thereby leading to elongated cells. Similarly, differences in exocellular properties between mutants would lead to a difference in how cells scatter light. Finally, an implicit assumption in our DMS analysis is that the allele frequency reflects cell frequency (i.e. differences would occur if the deleterious mutations caused selection for cells with fewer copies of the *TEM-1* plasmid). Regardless of the origin of the differences, the monoculture and co-culture growth experiments confirm that mutations identified as deleterious in the DMS experiment cause decreased growth rates.

### Comparison of primary and collateral fitness effects of mutations in TEM-1

We next examined the relationship between collateral and primary fitness effects in TEM-1. We calculated the primary fitness effects of TEM-1 missense mutations from data from an experiment by Stiffler et al (23). They performed a growth competition experiment on a comprehensive set of amino acid substitutions in the mature protein of TEM-1. They sought to quantify the effect of missense mutations on bacterial growth as a function of Amp concentration. At the start of their growth competition experiment, they divided a single culture into different cultures lacking or containing different concentrations of Amp. After a 10-fold increase in cell density, they compared the final allele frequencies in cultures containing Amp to the final allele frequency in the culture that lacked Amp, which they used to calculate an enrichment as their fitness metric. Under the assumption that collateral fitness effects are independent of Amp, this experiment measured the primary fitness effects of mutations (**Fig. 3A**). We used their sequencing count data to determine the primary fitness values based on mean growth rate (**Data S2-S5**), allowing for a direct comparison of primary and collateral fitness effects of mutations in TEM-1. As expected, deleterious primary fitness effects could be very large and frequent at high Amp concentration (**Fig. 3B**). The DFE of primary fitness effects was bimodal, with one peak centered around wildtype fitness and the other corresponding to lethal mutations. Primary fitness effects decreased in frequency and magnitude with decreasing Amp concentration (**Fig. 3B**,**C**). Thus, TEM-1 is buffered with respect to primary fitness effects of mutations unless the environment contains very high levels of Amp (23). Only mutations with drastic effects on protein activity/abundance cause detectable deleterious primary fitness effects in the low Amp environment.

**Figure 3.**
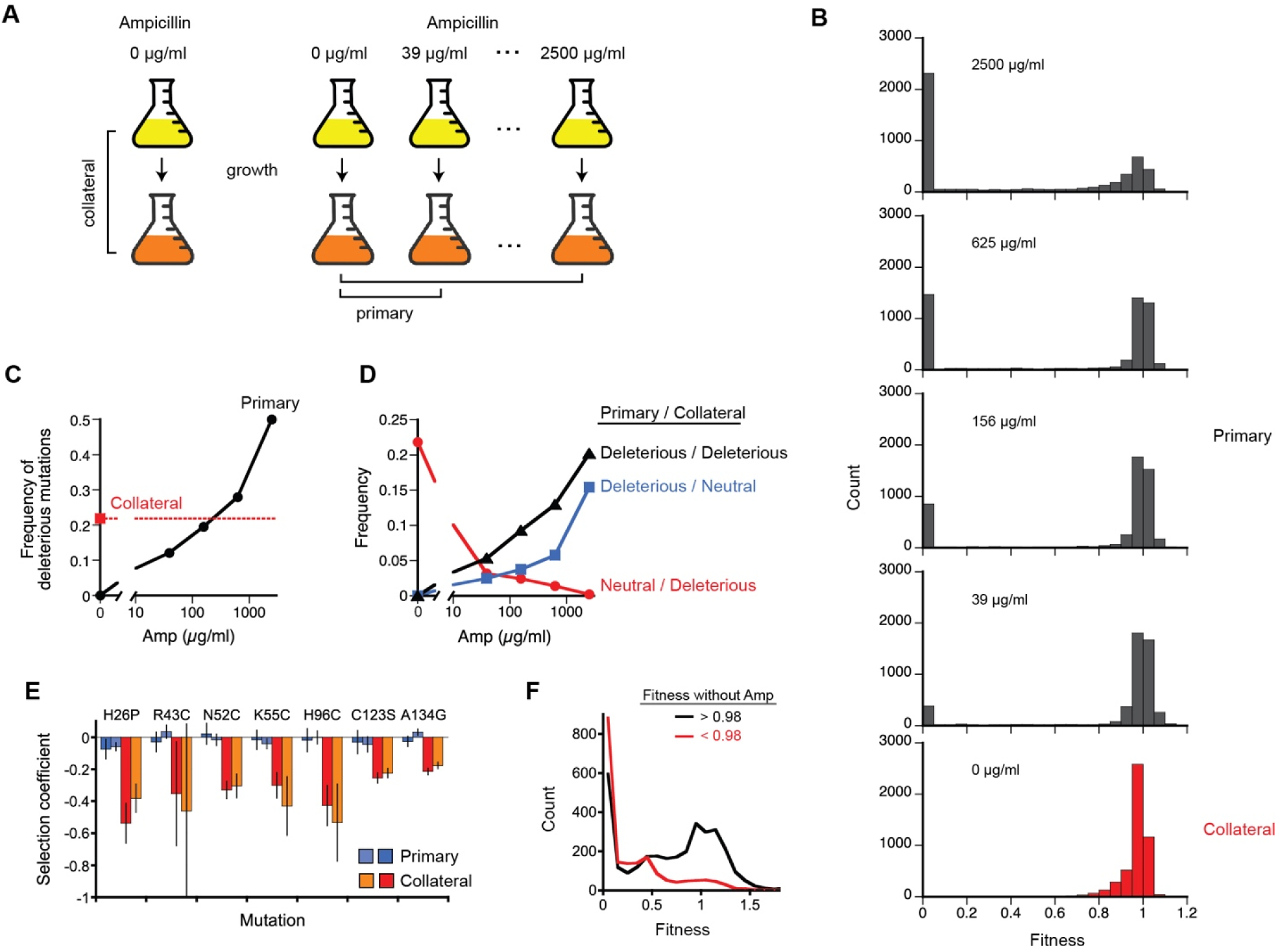
Comparison of primary and collateral fitness effects of mutations in TEM-1 β-lactamase. (**A**) Collateral fitness effects are measured by comparison of allele frequency before and after the growth period in the absence of Amp (left). Primary fitness effects are measured by comparison of the final allele frequency with and without Amp (right) in an experiment by Stiffler et al. (23). (**B**) Distribution of primary fitness effects (top four graphs) and collateral fitness effects (bottom graph) as a function of the indicated Amp concentration. For the primary fitness effect distributions, fitness measurements less than zero were set to zero. (**C**) Frequency of deleterious missense mutations with primary/collateral fitness effects in mature TEM-1. Criteria is *P*<0.01 in both replica experiments. (**D**) Frequency of missense mutations in mature TEM-1 with the indicated pattern of deleterious/neutral primary/collateral fitness effects. Neutral includes near-neutral mutations (see Methods). Heat maps of the mutations’ effects according to these categories are provided as **Fig. S10**. (**E**) Examples of mutations with deleterious collateral fitness effects and neutral/near-neutral primary fitness effects in media containing 156 µg/ml Amp. N52C, K55C and H96C also lack primary fitness effects at 625 µg/ml Amp (**Table S1**). (**F**) DFE of missense mutations in TEM-1 in the presence of Amp as measured by Firnberg et al. (31) divided into those mutations with deleterious collateral fitness effects (*w*<0.98; red) and those lacking deleterious collateral fitness effects (*w*>0.98; black).

We focused on deleterious fitness effects to examine the extent to which primary and collateral fitness effects contribute to the DFE in TEM-1. At the lowest Amp concentration, mutations caused collateral fitness effects more frequently than primary fitness effects (**Fig. 3C**). Primary fitness effects became as prevalent as collateral effects at intermediate Amp concentrations and were about twice as frequent at the highest Amp concentration. We then asked whether there was a correlation between mutations with primary and collateral fitness effects. For the purposes of this analysis, we classified each mutation as having only primary fitness effects, only collateral fitness effects, both primary and collateral, or the effects fit none of the three categories (**Fig. 3D, Fig S10**). This analysis showed that some mutations caused only collateral fitness effects, even at concentrations as high as 625 µg/ml Amp (**Fig. 3D, Table S1**). These mutations included several for which the collateral fitness effects were confirmed by growth assays (**Fig. 3E**). At the highest Amp concentration tested, which was just below the concentration that impaired growth of cells with the wildtype allele (23), 93% of mutations with deleterious collateral fitness effects also caused deleterious primary fitness effects (**Fig. 3D**).

We also compared the collateral fitness landscape to our previously described fitness landscape of the Amp resistance phenotype (31), which is predictive of the primary fitness effects in the presence of Amp (32). We found that mutations causing deleterious collateral fitness effects tended to be mutations that also decreased Amp resistance (**Fig. 3F**). Fitness in the presence of Amp was 5.6-fold lower when the collateral fitness effect was *s* < −0.02 than when it was not (*P* < 0.0001, Mann-Whitney; median fitness values were 0.142 and 0.791, respectively). This correlation would arise if deleterious primary and collateral fitness effects tend to have a common origin (e.g. in effects on protein folding/stability).

### Deleterious signal sequence mutations cause improper preTEM-1 processing, impaired release from the membrane, and aggregation

We next examined how the mutations affected the export, precursor processing, folding, and abundance of TEM-1. The pathway for exporting TEM-1 to the periplasm provides a framework for understanding potential mechanisms of deleterious signal sequence mutations. PreTEM-1 is exported to the periplasm by the Sec system via the SecYEG translocon. PreTEM-1 is fully synthesized prior to export (33). The Sec system exports proteins before they acquire tertiary structure. For Sec-exported proteins, a variety of molecular chaperones and the protein’s own signal sequence can help prevent acquisition of tertiary structure (34). Unlike most proteins exported by the Sec system, the cytoplasmic chaperone pair GroEL/GroES mediates the export of TEM-1 (35) by binding to and stabilizing the unfolded state (36). Although the mature domain of preTEM-1 can fold into an active enzyme without removal of the signal sequence (37), preTEM-1’s signal sequence stabilizes the unfolded state, and preTEM-1 folds 15-times slower than TEM-1 (37). All Sec pathway proteins interact with SecA, which interacts with the hydrophobic portion of the signal sequence. A Sec signal sequence is characterized by a positively charged N-terminal region, a hydrophobic core region, and a neutral C-terminal region that contains the cleavage site immediately after a consensus A-X-A motif (often preceded by a helix breaking residue, typically a proline, 1-3 residues away) (38) (**Fig S11A**). The last four residues of TEM-1’s signal sequence (residues 22-25) are PVFA, with a valine replacing the first alanine in the A-X-A motif. Post-translocational cleavage of TEM-1’s signal peptide by signal peptidase I (SPaseI) catalyzes the release of mature TEM-1 to the periplasm (39).

We observed deleterious collateral fitness effects with *s* < −0.05 at 7 of the 23 positions in TEM-1’s signal peptide and when Pro was substituted for the first amino acid of the mature protein (**Fig S11B**). The most frequent and most deleterious effects occurred at P22 and A25, suggesting improper cleavage of the signal peptide might cause many of the deleterious effects. We chose twelve mutants to study, representing most of the positions displaying fitness effects. Cell fractionation experiments and subsequent western blot analysis indicate that all twelve mutations caused a defect in preTEM-1 processing resulting in precursor aggregation in the inclusion body fraction (**Fig. 4A)**. Failure to remove the signal sequence would favor aggregation since the signal sequence reduces TEM-1’s folding rate 15-fold (37). Some mutations compromised release of the precursor from the membrane. These processing defects manifested in different ways depending on the site of mutation. Mutations at P22 uniquely caused a species intermediate between the precursor and the mature protein to accumulate in the insoluble and membrane fraction (**Fig. 4D**). Mutations at L12 and V23 caused TEM-1 molecules of approximately the same size as the mature protein to aggregate along with preTEM-1. Several mutations, typically the most deleterious, prevented mature protein formation.

**Figure 4.**
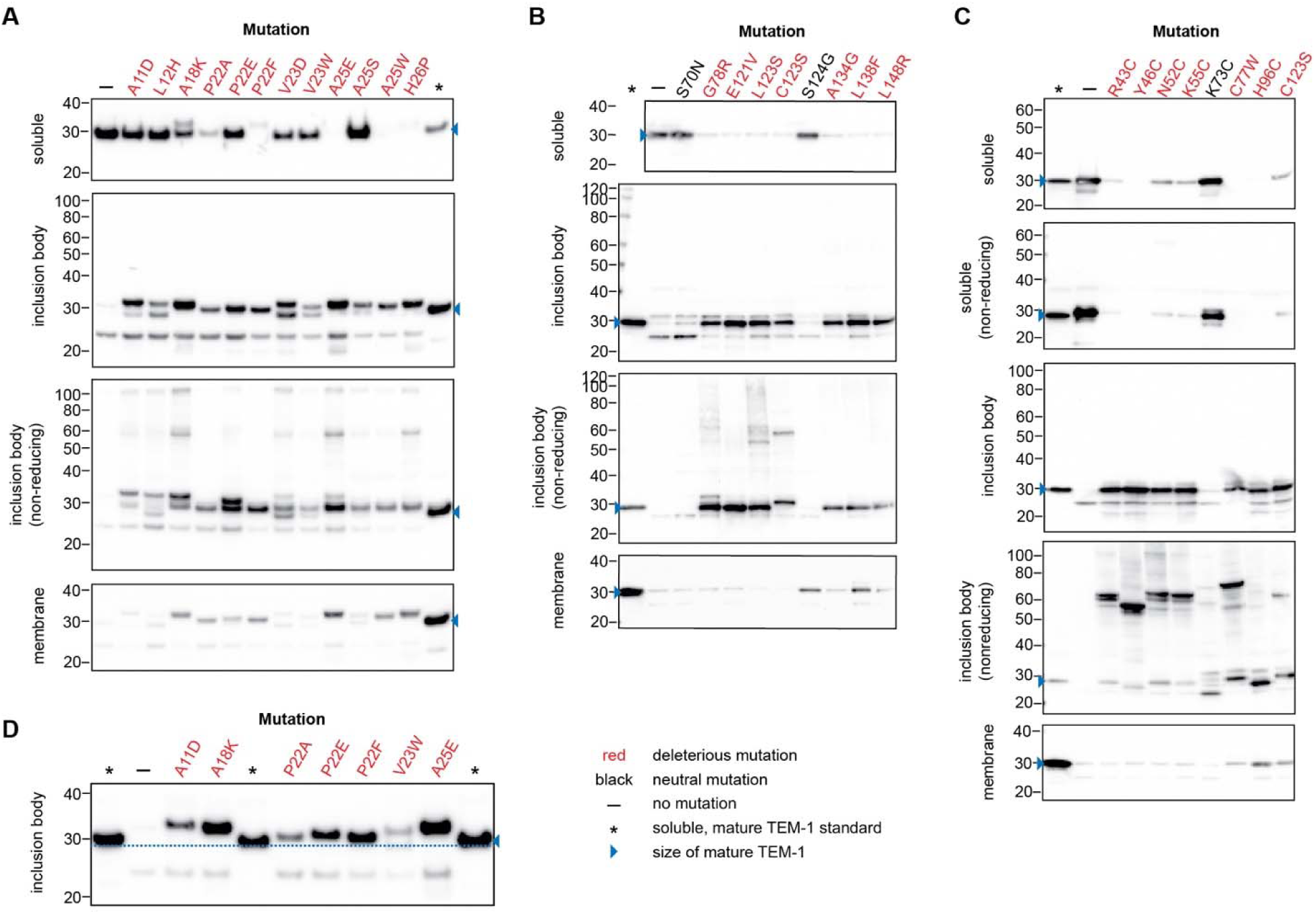
Effects of mutations on TEM-1 expression, processing, folding, and disulfide bond formation. Exponential phase cultures expressing TEM-1 with the indicated mutations were harvested after six hours of exponential growth (post-induction). Cells were fractionated into soluble (periplasmic or total), inclusion body, and membrane fractions; separated by reducing SDS-PAGE (or non-reducing, as indicated); and probed by western blot with polyclonal anti-TEM-1 antisera. Mutations tested included those (**A**) in the signal sequence, (**B**) in α-helices of the α domain, and (**C**) involving cysteine residues. (**D**) is a western blot illustrating that select signal sequence mutations cause improper signal sequence processing. The blue dashed line is provided at the bottom of the band of the mature TEM-1 as a guide for the eye. The soluble fraction in (A, C) is the periplasmic fraction. The soluble fraction in (B) is the total soluble fraction. Inclusion body and membrane samples are loaded at a higher level per OD unit than soluble samples in order to better see the mutations’ effects. Deleterious mutations are labeled in red. Neutral mutations are labeled in black. The blue arrowhead indicates the migration position of mature TEM-1. Full western blots and western blots of samples from replicate cultures provided as **Figs. S14, S15, and S16**.

In the hydrophobic core of the signal sequence, the introduction of charged residues at A11, His at L12, and positively charged mutations at A18 caused collateral fitness effects and a defect in preTEM-1 processing (**Fig 4A**). Mutations in the hydrophobic core have not been previously linked to deficiencies in signal sequence cleavage and release from the inner membrane. Such deficiencies have been limited to mutations in the C-terminus of the signal sequence such as P22S, P22F, P22L and A25E (40-42). The introduction of charged residues at most positions in the hydrophobic core causes a large decrease in the ability of the gene to provide Amp resistance (31), consistent with typical export-defective mutants in Sec pathway signal sequences (43). However, we did not observe a collateral fitness cost for most of these mutations, suggesting that disruption of TEM-1 export does not have a fitness cost per se. Perhaps defective preTEM-1 processing (rather than a decrease in export efficiency) leads to collateral fitness effects since all tested deleterious mutations in the signal sequence cause a processing defect. Our cell fractionation experiments do not distinguish whether the insoluble preTEM-1 is in the periplasm or the cytoplasm. However, preTEM-1 aggregation is most likely in the periplasm for deleterious mutations near the signal peptide C-terminus because this region is not required for transport across the membrane but determines the rate of signal sequence cleavage and release from the membrane (40). For deleterious mutations in the hydrophobic core region, the mutations might disrupt interactions with cytoplasmic chaperones or SecA, compromising export and resulting in cytoplasmic preTEM-1 aggregation.

### Deleterious mutations in α-helices of the α-domain cause aggregation of mature TEM-1

In vitro folding of TEM-1 occurs via intermediates in which the α domain is collapsed and partially folded (i.e. molten globule like), and the two lobes of the α/β domain have not associated (44). The C-terminal region has transient non-native interactions in these intermediates (45). Formation of native TEM-1 occurs via two major parallel pathways, both involving the rate-limiting step of trans to cis isomerization of the E166-P167 bond in the Ω-loop (46), in which the N- and C-terminal lobes of the α/β domain associate. Previous neutral drift mutation accumulation experiments enriched for stabilizing mutations in TEM-1’s α domain (R122G, E149G, H155R, M184T, and L203P) (47). These mutations affected the folding intermediates by providing thermodynamic and/or kinetic stability, which suggested that the outcome of TEM-1 folding depends on the starting configuration of the α domain. In our experiment, all five stabilizing mutations were neutral. In contrast, many mutations to the α domain, especially in α-helices 4 and 5, caused deleterious collateral fitness effects (**Fig. 1A,D**). We wondered if these deleterious mutations had the opposite effect of the stabilizing mutations.

We chose to mutate positions in TEM-1 in α-helices 2, 4, 5, and 6 at which most mutations had negative effects (**Fig S13**). In this set of mutants, we introduced single-bp substitutions to sample exclusively from *TEM-1*’s nearest neighbors – the type of mutations most likely to occur naturally. We chose S70N and S124G as negative control mutations that lacked collateral fitness effects. Cells expressing TEM-1 with deleterious helix mutations produced little if any soluble TEM-1; instead the protein aggregated (**Fig. 4B**). These results reinforce the α domain’s importance in TEM-1’s folding pathway. We postulate that the mutations perturb the state of the α domain in TEM-1 folding intermediates causing off-pathway aggregation to occur in the periplasm.

### Deleterious mutations involving cysteine cause improper intermolecular disulfides and aggregation

TEM-1 contains three cysteines: C20 in the signal sequence and C77/C123, which form a disulfide bond after TEM-1 is exported to the periplasm. The disulfide bond is not required for TEM-1 folding or catalytic activity (37). We observed deleterious fitness effects at all three cysteines, but especially at C123 (**Fig. S11B and S12B**). Elsewhere in the protein, mutations to cysteine tended to be more deleterious than mutations to other amino acids (**Fig. S6A**). Mutations to cysteine tended to be more deleterious than mutations to serine despite the two amino acids being polar molecules of equivalent size (**Fig. S12A**). This trend suggested that cysteine’s reactivity plays a role in its deleterious fitness effects. We chose to study the R43C, Y46C, N52C, K55C, and H96C mutations because cysteine uniquely caused deleterious fitness effects at these positions (**Fig S12B**). We also chose C77W and C123S as mutations at the native disulfide-forming cysteines. N52, K55, and H96 are partially exposed to solvent; Y46, C77, and C123 are completely buried (**Fig S12B, Table S1**). We chose K73C as a mutation with no deleterious collateral fitness effect.

Cells expressing TEM-1 with these deleterious mutations produced little if any soluble TEM-1 (**Fig. 4C**). Instead, the protein accumulated in inclusion bodies containing incorrect, intermolecular disulfides. All deleterious mutations involving cysteine except H96C caused the appearance of a species at about twice the molecular weight of TEM-1 (∼58 kDa), suggesting disulfide-linked TEM-1 dimers. Such a species occurred in a few mutants not involving cysteine but only as a minor species (**Fig. 4A,B**). The intermolecular disulfides formed in the cell (and not after lysis), as the same higher molecular weight species occurred when we added iodoacetamide to the cells prior to cell lysis to quench free thiols (**Fig S15**).

Whether the intermolecular disulfides are the cause or a consequence of aggregation is unknown, but we suspect the former. Many mutations to cysteine throughout the α domain and the N-terminal lobe of the α/β domain were more deleterious than serine, but the C-terminal lobe of the α/β domain lacked such mutations (**Fig S12A**). Since TEM-1 is secreted from N- to C-terminus, we speculate that cysteines introduced before the end of the α domain are prone to form intermolecular disulfides before the α domain is fully secreted and can rapidly collapse to its initial compact state. Formation of intermolecular disulfides would likely impair folding and lead to aggregation. However, after the α domain is secreted, α domain collapse might make formation of intermolecular disulfides less likely.

### Deleterious nonsense mutations correlate with the accumulation of a TEM-1 protein fragment

We examined the consequences of nonsense mutations in the deleterious-prone region between residues 170 and 195. Nonsense mutations designated as deleterious by DMS caused a particular pattern of protein expression. Deleterious mutations caused the appearance of two TEM-1 fragments in the inclusion body fraction, and the soluble fraction had a fragment at a size similar to that of the smaller fragment (**Fig. S17**). Non-deleterious mutations lacked the lower size fragment in both the soluble and insoluble fractions despite the location of the nonsense codon differing by only one position in some cases. Since the smaller insoluble fragment increased in size as the nonsense mutation moved further from the 5’ end of the gene, the fragment is likely produced by proteolytic cleavage near the N-terminus, perhaps by SPaseI cleavage of the signal sequence. This circumstantial evidence suggests that the smaller species has a role in the deleterious fitness effects.

### Deleterious mutations activate select stress-response pathways

We performed RNA-Seq experiments on a subset of mutants to learn how the mutations affected the cell’s gene expression profile. We identified genes and pathways of interest from those in which a mutation caused several genes in a pathway to have a >2-fold difference in expression and the *P* value was less than 0.001. Induction of TEM-1 expression caused virtually no change in gene expression (**Fig. S1D**) and no induction of stress-response pathways despite TEM-1’s moderately high level of expression. The addition of IPTG caused little if any fitness effect (**Fig. S2**) and little change in gene expression except for the expected upregulation of the *lac* operon (**Fig. S1E**). In contrast, deleterious mutations caused significant changes in gene expression (**Figs. S18**) primarily in the induction of specific outer envelope stress response pathways and, in some cases, the mild-induction of a few genes in the heat-shock response pathway (**Fig. 5**). The mutations did not all induce the same stress pathways, suggesting different origins for the deleterious fitness effects or differences in the cell’s ability to respond to the mutations’ effects. Although deleterious mutations in essential, core metabolic enzymes can produce widespread pleiotropic effects in gene expression (48), the effects we observe here, though less extensive, are for an enzyme whose catalytic activity is irrelevant for fitness.

**Figure 5.**
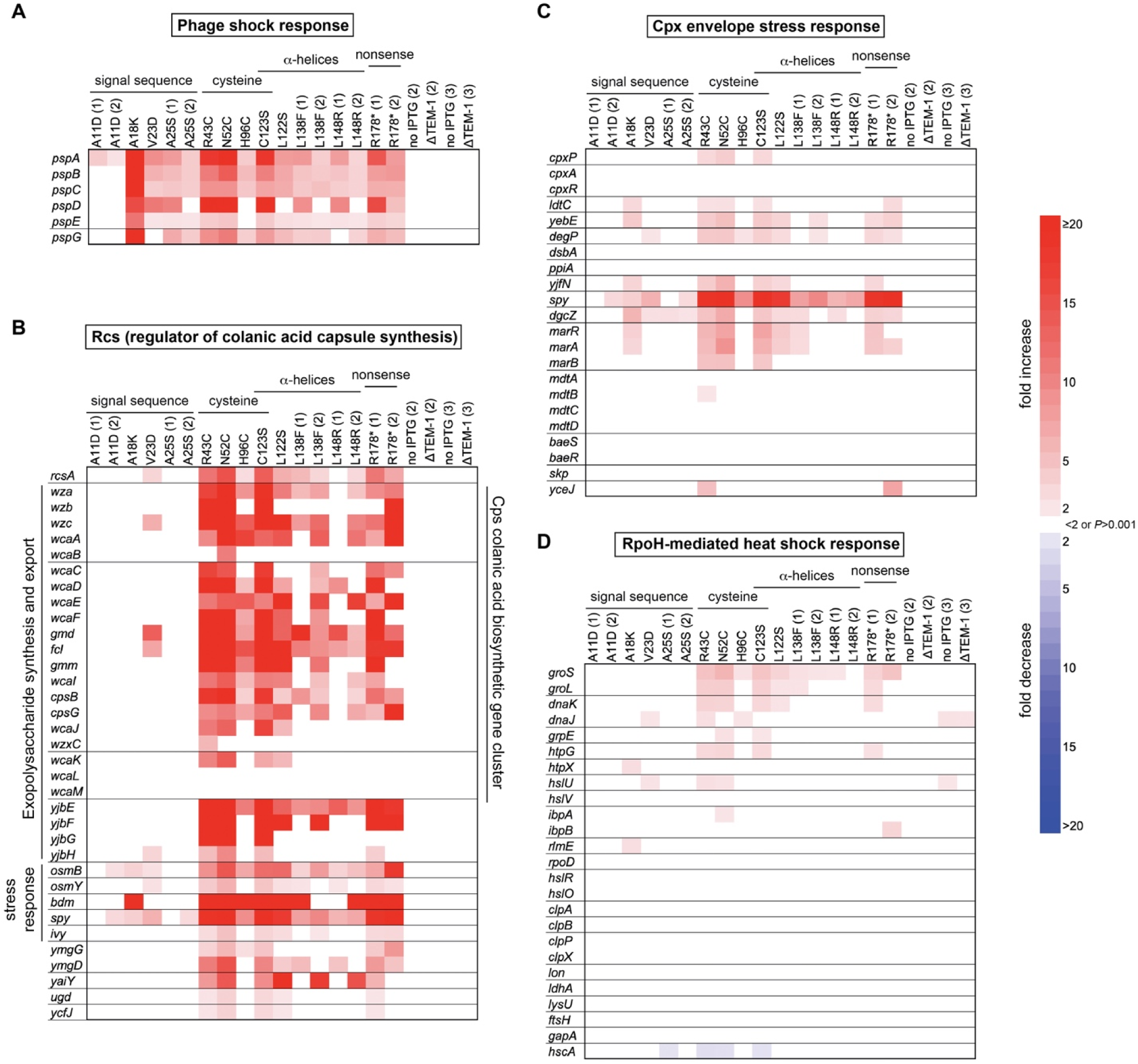
Effect of deleterious mutations on *E. coli’s* gene expression profile. RNA-Seq experiments were performed on exponentially growing cells 5.5 hours post-induction of TEM-1. The heatmaps show changes in gene expression (relative to cells expressing unmutated TEM-1) that differ by >2-fold (*P* < 0.001, *Z*-test) for select genes in the (**A**) phage shock protein pathway, (**B**) Rcs stress response pathway, (**C**) Cpx stress response pathway, and (**D**), σ^H^-mediated heat shock response pathway. The number in parentheses after the mutation is for samples with replica experiments and indicates the day of the experiment. The control experiments in the last four columns correspond to cells lacking induction of TEM-1 expression (no IPTG) or cells lacking the TEM-1 gene but induced with IPTG (ΔTEM-1). Additional observations in differential gene expression are provided as **Supplemental Text**. Data provided as **Data S6**.

The phage shock protein response (Psp) and the regulator of colanic acid capsule synthesis pathway (Rcs) were the most commonly-activated pathways and had the largest response. All but the A11D mutation (the least deleterious mutation tested) induced the two known Psp operons, *pspABCDE* and *pspG* (**Fig. 5A**). The Psp response is believed to be induced by stress on the inner membrane and mitigates problems that could increase inner membrane permeability (49). Signal sequence mutations affected Psp pathway gene expression more than they affected genes in other stress pathways. The largest set of genes upregulated were those in the Rcs stress response pathway (**Fig. 5B**). The known triggers of the Rcs stress response pathway are outer membrane or peptidoglycan layer stresses caused by drugs and the switch from planktonic, motile growth to nonmotile growth and biofilm formation (50). Protein aggregation has not been described as an Rcs trigger. Of the mutations studied here, all mutations to the mature protein led to upregulation of genes controlled by the Rcs pathway while signal sequence mutations generally did not. In particular, A18K and A25S caused almost no detectable upregulation of genes in this pathway despite being the most deleterious signal sequence mutations tested. The set of upregulated Rcs genes included the large *cps* gene cluster responsible for production and secretion of the exopolysaccharide (EPS) colanic acid (51), the *yjbEFGH* operon (implicated in the production of a different EPS (52)), and several stress response genes including *ivy, osmB*, and the protein misfolding chaperone *spy.*

Spy is also known to be induced in the Cpx envelope stress response pathway. The Cpx response is thought to be primarily tasked with maintaining inner membrane integrity (53, 54). It is stimulated by misfolded periplasmic and inner membrane proteins (but not misfolded outer membrane proteins) and by inner membrane stress including translocation defects. In addition to increasing *spy* expression, select *TEM-1* mutations increased the expression of the periplasmic protease *degP* and its activator *yjfN*, which are part of the Cpx regulon (**Fig. 5C**). However, many genes in the Cpx response did not exhibit differential expression, and for those that did, the changes were generally small.

Genes in the two remaining major outer envelope stress response pathways, the σ^E^ and BaeR stress responses, did not show differential gene expression. The σ^E^ stress response is stimulated by misfolding of outer membrane proteins (55), and the BaeR stress response is stimulated by periplasmic protein misfolding (56). The most deleterious mutations in the mature protein mildly stimulated expression of a few genes in the σ^H^-mediated heat-shock response (a stress response that is also stimulated by cytoplasmic protein misfolding) (**Fig. 5D**). These genes included cytoplasmic chaperone *groS/groL*, which is known to interact with unfolded preTEM-1 (35, 36). However, the expression increase was low compared to that typical of the heat shock response, and many heat-shock genes did not show a significant change in gene expression. We rarely observed any differential expression for genes in the σ^S^-mediated general stress response, which responds to a large variety of stresses including temperature, osmotic shock, pH, and oxidative stress (57). The relatively mild overexpression of periplasmic/cytoplasmic chaperones (as a whole) is surprising given all mutations studied caused TEM-1 aggregation.

### Deleterious mutations cause pleiotropic changes in cell phenotype

Pleiotropy complicates the evolution of proteins. The mutations studied here have a variety of effects on the β-lactam antibiotic resistance phenotype (**Table S1**) but also changed the expression level of many genes (**Fig. 5**), which might cause other phenotypic changes. Since many of the differentially expressed genes involve the outer envelope of the cell, we wondered if deleterious mutations cause any changes to the ability of the cells to interact with their environment and, in particular, changes that would produce a visible phenotypic change.

As an example, we examined a phenotype associated with EPS production. One of the strongest transcriptional responses to many of the deleterious mutations was the upregulation of the Rcs regulon involved in the production and transport of EPS. We used a dye uptake assay to assess whether the mutations caused an increase in EPS. Increased expression of the *yjbEFGH* operon causes *E. coli* colonies to bind Congo red (known to recognize (1→4)-α-D-glucopyranosides and basic or neutral EPS) and toluidine blue-O (known to recognize negatively charge or helical polysaccharides) (52). We found that only mutants with a strongly activated Rcs response caused Congo red uptake (**Fig. 6A,B**). Toluidine blue-O uptake varied but occurred with some of the most deleterious mutations (**Fig. 6C, Fig. S20CD**). All deleterious mutants in mature TEM-1 caused Congo red uptake, but in the signal sequence, only the most deleterious mutants at P22 and A25 caused dye uptake (**Fig. S20AB**). Nonsense mutations causing Congo red uptake were limited to those with deleterious fitness effects as measured by DMS or steady-state growth rate, corroborating these mutants’ effects. Cells expressing the most deleterious mutants formed smaller colonies (**Fig. 6BC, Fig. S20AC**), illustrating that deleterious mutations caused fitness defects on solid media. These results illustrate that mutations causing collateral fitness effects can also change how the cell interacts with its environment, which may have important consequences for the fitness landscape in some environments.

**Figure 6.**
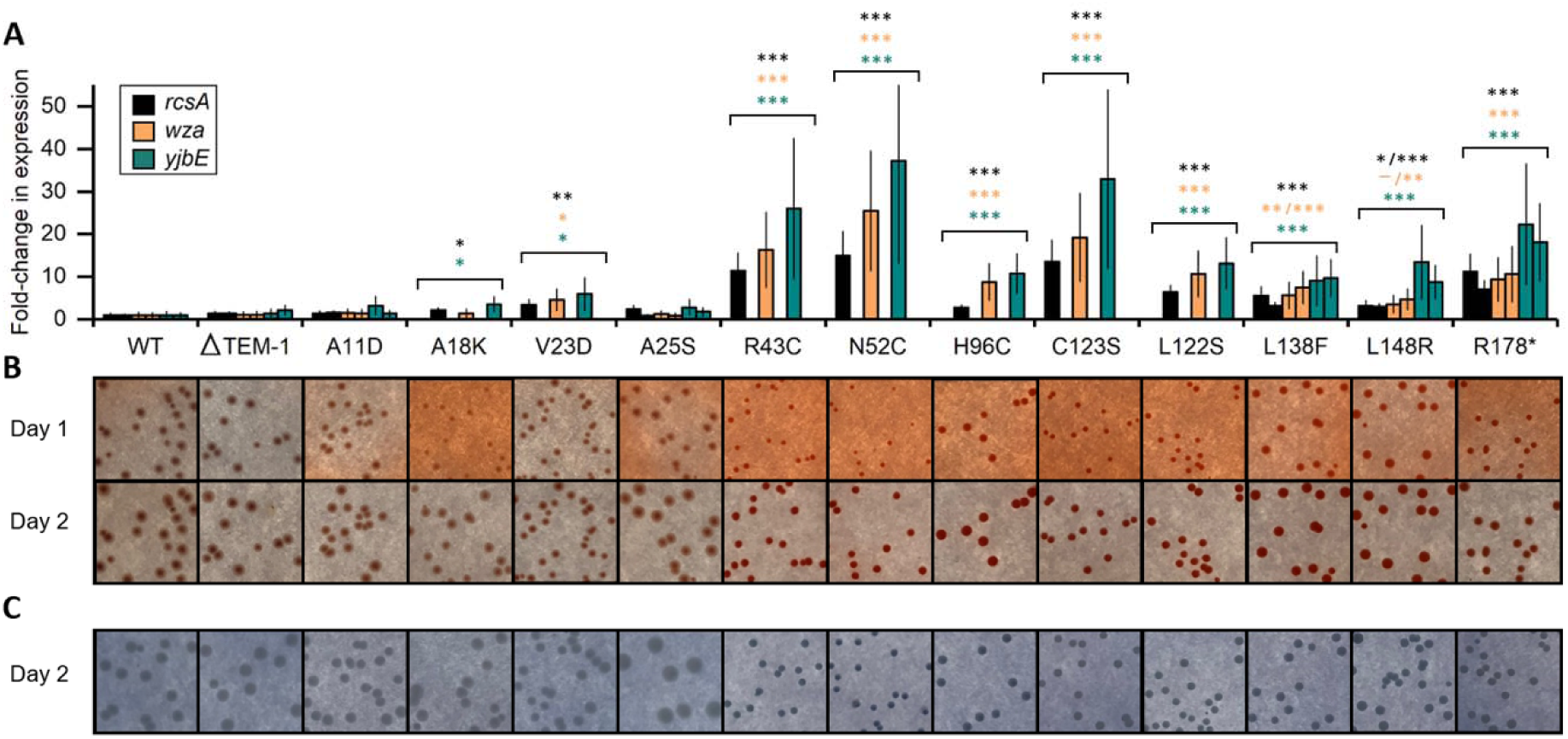
Pleiotropic effects of deleterious mutations. Cells expressing wildtype and mutant proteins were assessed for changes in gene expression and dye uptake related to Rcs-regulated EPS production on the outside of *E. coli* cells. (**A**) Differential gene expression of *rcsA* (accessory regulator of Rcs pathway) and two genes induced by the Rcs signal cascade: *wza* (first gene in the *cps* gene cluster for producing the EPS colanic acid), and *yjbE* (first gene in the *yjbEFGH* operon implicated in EPS production). Mutants with multiple bars represent replica experiments. Error bars are the standard deviations. Mutations that cause significant changes in gene expression are marked with asterisks colored according to the corresponding gene (*, *P*<0.01; **, *P*<0.001; ***, *P*<0.0001, *Z*-test). (**B**,**C**) Induced, exponentially growing cultures were diluted and spread on LB plates supplemented with IPTG and either (B) Congo red or (C) toluidine blue-O. Plates were incubated for two days at 37°C. Dye uptake indicates EPS production. Day 1 images show the mutations’ effect on colony size. On day two, colonies for R43C and those mutants to the right of R43C are more red/blue than the colonies to left of R43C. Dye uptake of additional mutants provided as **Fig. S20**.

## Discussion

Our study establishes that missense mutations in *TEM-1* often cause deleterious collateral fitness effects, at least at the expression level studied here. Our results caution against using measures of protein properties as a proxy for biological fitness or employing a non-native context in DMS experiments, as the former risks underestimating the fraction of mutations that are deleterious in their native context and the latter risks reporting on non-biologically-relevant collateral fitness effects. The collateral fitness effects we observed were associated with TEM-1/preTEM-1 aggregation, improper signal sequence cleavage, impaired release of the mature protein from the membrane, incorrect disulfide-bond formation, induction of stress response pathways, and pleiotropic changes in cell phenotype. Our study does not establish the relationship between these observations, or which, if any, cause the fitness decrease. A circumstantial case can be made for aggregation being the root cause of the deleterious effects. All 25 deleterious missense mutations caused aggregation, but none of the three neutral mutations did.

If aggregation is the cause of the deleterious effects, why are stress response pathways known to be stimulated by protein aggregation only weakly activated and only for some mutants? We postulate that because misfolded proteins are not toxic in vacuo (i.e. they must interact with some cellular component to cause deleterious fitness effects), the cell’s response to the aggregates depends on the nature of the misfolded protein (which may be mutation-dependent), how the misfolded protein interacts with some cellular component to cause a deleterious fitness effect, and whether a stress response can detect and respond to the consequences of that interaction. Thus, the strong activation of the Psp and Rcs pathways (whose activation has not been previously attributed to protein misfolding per se) could result from misfolded TEM-1 perturbing cellular processes or molecules that are monitored by these stress response systems. However, we did not examine whether the mutations caused either misfolded or correctly folded TEM-1 to misinteract with other proteins. In addition, fitness costs could arise from the inadvertent induction of a stress response when none is warranted and from the cost of removing the aggregated protein. In terms of a mutation’s impact on protein evolution, the cause of the deleterious effects is irrelevant. Collateral fitness costs arise from the mutation’s impact on some cellular process(es) that in turn causes a fitness decrease. These costs will constrain protein evolution, provided they are not offset by beneficial primary fitness effects or mitigated by a second, compensatory mutation that alleviates the effect or lowers expression level.

Under what conditions will collateral fitness effects be important to protein evolution? A gene’s contribution to fitness may be environment-dependent (e.g. *TEM-1* is essential for growth only in the presence of β-lactam antibiotics). In an environment where the gene is expressed and not under selection for its physiological function, deleterious collateral fitness effects will constrain its evolution (**Fig. 7A**). For genes under selection, such as *TEM-1* in environments with β-lactam antibiotics, small-effect mutations on the protein’s specific activity or abundance may be neutral on the level of fitness due to the non-linear mapping of protein physicochemical properties on organismal fitness (11-13) (**Fig. 7B**). This non-linear mapping occurs with respect to protein abundance because of the threshold behavior of the Boltzmann distribution that describes how the probability of the folded state depends on the free energy of folding (14), and because protein chaperones act as a buffer for deleterious mutational effects on soluble protein abundance (58, 59). Buffering can also occur with respect to specific protein activity. If an essential metabolic pathway is more than sufficient for producing a metabolite, which is often the case, then mutations causing small changes in the metabolic enzyme’s catalytic activity may be selectively neutral, at least as far as the protein’s physiological function is concerned (10, 60). For example, mutations in the essential *E. coli* enzyme dihydrofolate reductase that span a 14-fold range in catalytic activity, a 5-fold range in protein abundance, and a 28-fold range in total cellular catalytic activity had fitness values that spanned only a 1.8-fold range, and these values did not correlate with total cellular activity (11). Genes whose primary fitness effects are buffered against small-effect changes in specific activity or abundance will remain vulnerable to deleterious collateral fitness effects (**Fig. 7B**), which will constrain their evolution.

**Figure 7.**
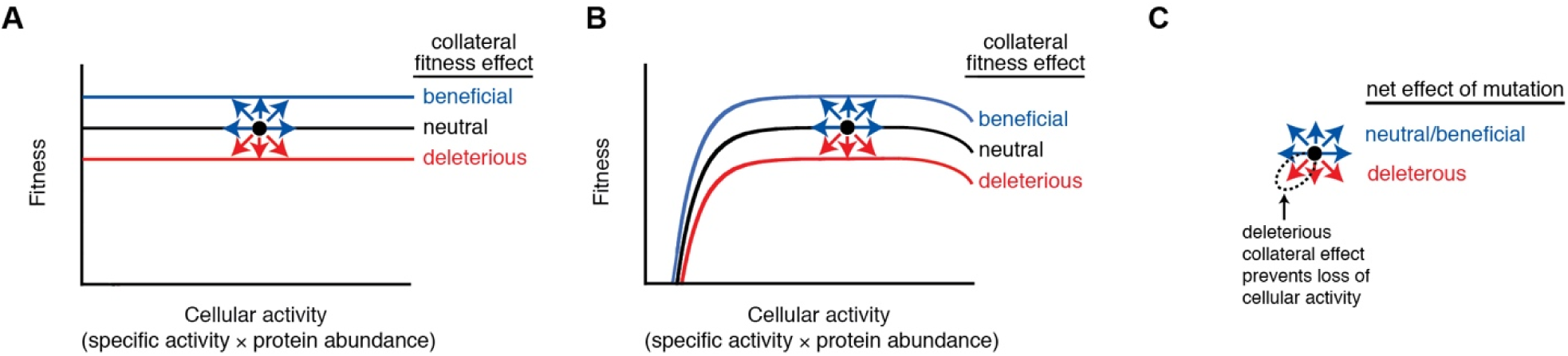
Evolutionary consequences of collateral fitness effects. The model is illustrated by hypothetical relationships between fitness and cellular activity for an enzyme that has a singular physiological function. The enzyme is either (**A**) not under selection or (**B**) under selection for its physiological function. Genes under selection for their physiological function have diminished fitness if they provide too little or too much cellular activity. At optimal values of cellular activity, fitness is buffered against changes in cellular activity above a threshold amount. Black circles represent an enzyme subject to mutation. A mutation having an effect on cellular activity moves the protein along a line. A mutation having a collateral fitness effects moves the protein vertically to new lines. (**C**) The net effects of mutation on the protein (arrows) are colored by whether they are deleterious and will be purged by evolution (red) or neutral/beneficial and offer possible evolution pathways (blue). The downward left arrow represents mutations that causes both deleterious collateral effects on fitness and deleterious effects on cellular activity. For such mutations, the deleterious collateral fitness effects can prevent the protein from acquiring mutations that compromise cellular activity if one of the following is true: 1) the enzyme is not under selection for its physiological function or 2) the gene is under selection for its physiological function and the deleterious effect on cellular activity is not large enough to move the enzyme from the buffered region. Collateral fitness effects will dominate the constraint on evolution in this buffered region. We did not conclusively establish that mutations can cause a beneficial collateral fitness effect in TEM-1; the possibility of such effects are included in this general model.

Our data suggests that mutations with deleterious collateral fitness effects are biased to protect against genetic drift that would compromise the gene’s physiological function. This phenomena arises because mutations causing collateral fitness effects tend to also cause primary fitness effects (**Fig. 3D&F, Fig. S10**). In some environments, mutations might only have collateral fitness effects, either because the protein is not under selection (**Fig. 7A**) or because primary fitness effects are buffered against mutational effects on cellular activity (**Fig 7B**). Thus, in such environments, deleterious collateral fitness effects decrease the likelihood of the gene accumulating missense mutations that would compromise its physiological function – mutations that would erode the primary fitness effect buffer or be detrimental to fitness if the gene were to later come under selection for its physiological function (**Fig. 7C**). We postulate that deleterious collateral fitness effects help maintain gene function and preserve mutational robustness in genes that are intermittently under selection. Such a phenomenon would depend on the time scale of environmental changes and the frequency of mutation. One might imagine that when a constitutively-expressed gene is not under selection for its physiological function, mutations that decrease or eliminate expression will be selected (e.g. through a promoter mutation or a nonsense mutation early in the gene), a scenario which would seem to reduce the evolutionary importance of collateral fitness effects. However, it is the fitness across all environments an organism experiences that will govern the gene’s evolution. When the gene comes back under selection, such silenced alleles will be rapidly purged because they are not functional. Environment-dependent regulation of gene expression could alleviate this issue; thus, collateral fitness effects might provide an evolutionary advantage for the evolution of gene expression regulation.

If collateral fitness effects primarily arise from protein aggregation/misinteraction, then collateral fitness effects’ constraint on protein evolution would increase with expression level because aggregation and interactions are concentration-dependent. Thus, our study is potentially relevant to the observation that the best predictor of protein evolution rates is an anticorrelation with protein expression level. This phenomenon is called the E-R anticorrelation, where E stands for gene expression and R stands for evolutionary rate. The reasons that proteins evolve at different rates has been a longstanding debate in evolutionary and molecular biology (61, 62). Protein evolution rates are critical to the reconstruction of evolutionary history and mechanisms. According to the neutral theory of evolution, the rate of protein evolution equals the rate of mutation multiplied by the proportion of mutations that are neutral (63). The proportion of neutral mutations is set by functional constraint, which is the extent to which mutations are purged by natural selection due to their deleterious effects. Interestingly, the functional importance or essentiality of a protein is not the major determinant of protein evolutionary rates (62). Although many correlates of protein evolution rates have been noted (62), the best predictor is protein expression level, which is anticorrelated (3, 4, 7, 64). Major hypotheses for the E-R anticorrelation posit that mutant proteins tend to misfold and misinteract. Misinteractions can originate from a protein either in its native state (3-6) or a misfolded conformation (7-9) so as to cause deleterious effects on fitness. If misfolding/misinteractions are an important contributor to the E-R anticorrelation, deleterious collateral fitness effects should be frequent in a highly expressed protein such as TEM-1. Our observation that at least 42% of mutations have deleterious collateral fitness effects is consistent with this expectation.

TEM-1 was an excellent protein for studying collateral fitness effects because its high expression increased the chance for consequential misfolding/misinteractions and its export allowed for the observation of fitness effects arising from several different mechanisms (i.e. effects in two cellular compartments and those related to export). Are the frequency and magnitude of collateral fitness effects of mutations in TEM-1 representative of other *E. coli* proteins? Are periplasmic proteins more prone to collateral fitness effects than cytoplasmic proteins due to the need for export across the inner membrane and the potential for misinteractions in two cellular compartments? These questions can only be answered by similar studies on other genes. The difficulty in performing such studies comes in assessing whether or not the measured fitness effects arise independent from their effects on the protein’s physiological function. In addition, our study considered the fitness effects in a single environment and a single phase of growth. Fitness effects of mutations (even the sign of the effect) can vary greatly with environment (20) and may be growth phase dependent, which has important consequences for selection (65). A key assumption of our study is that TEM-1’s known catalytic activity (or any unknown secondary activity) does not affect any biological process in the cell that impacts fitness. We expect that *TEM-1’s* plasmid-borne nature and seemingly singular, conditional purpose make it less likely to participate or impact other cellular processes. The lack of fitness effects or gene expression changes upon *TEM-1* expression supports this position. Our study indicates that collateral fitness effects are potentially frequent for highly expressed genes, which is a requirement for the misfolding and misinteraction hypotheses for the E-R anticorrelation, and raises the possibility that protein sequence evolution is under selective constraint both to maintain function and to avoid collateral negative effects.

## Materials and Methods

Details of the materials and methods, including library construction, growth competition experiments, DMS and data analysis, fitness measurement methods, cell fractionation analysis, RNA-Seq and data analysis, and data availability are presented in **Supplementary Materials and Methods**.

## Acknowledgements

We thank M. A. Stiffler for providing the sequencing count data from (23). We thank C. Kaiser, D. Tawfik, and W. Patrick for comments on a draft version of the manuscript. This research was supported by National Science Foundation grants DEB-1353143 and MCB-1817646 to M.O. A.F.R benefited from an Australian National Health and Medical Research Council (NHMRC) Program Grant (1054618). The research benefited by support from the Victorian State Government Operational Infrastructure Support and Australian Government NHMRC Independent Research Institute Infrastructure Support.

## Author contributions

J.D.M. Obtained collateral fitness data on the last third of the gene, did the final analysis of all DMS sequencing data using Enrich2, constructed and characterized the α domain mutants, analyzed the sequencing count data from Stiffler et al. (23) to calculate primary fitness effects, performed the RNA-Seq experiments and analysis, confirmed the fitness effects of mutations in co-culture, performed dye plate experiments, supervised some experiments done by co-authors, wrote the methods section, and edited the manuscript. F.W.S did several preliminary experiments, obtained collateral fitness data for the first two-thirds of the gene, and measured fitness effects of Cys mutants in monoculture. D.R. constructed and characterized the Cys mutants and edited the manuscript. B.W. constructed and characterized the signal sequence mutations. E.-Y.T. constructed and characterized the nonsense mutations and performed dye plate experiments. N.D. characterized the α domain mutants. M.-H.H. measured the fitness effect of TEM-1 induction and constructed and measured the fitness effects of the synonymous mutants. C.E.G. provided initial analysis of the DMS sequencing data using MATLAB scripts. A.F.R. provided preliminary analysis of the DMS sequencing data using Enrich2, assisted with the Enrich2 software that included adjustments to suit this project, and edited the manuscript. M.O. conceived of the study, acquired funding, supervised the work, and wrote the manuscript. All authors analyzed data, discussed the results, and reviewed the manuscript.

