## Supplementary Materials for "Collateral fitness effects of mutations"

**This PDF file includes:**

Supplementary Materials and Methods
Supplementary Text
Figs. S1 to S20
Table S1
Supplementary References
Captions for Data S1 to S6

**Other Supplementary Materials for this manuscript include the following:**

Data S1 to S6

### Materials and Methods

**General methods.** The strain was NEB 5-alpha LacI$^q$ (F´ *proA*$^+$*B*$^+$ *lacI*$^q$ *Δ(lacZ)M15 zzf::Tn10* (Tet$^R$) */ fhuA2Δ(argF-lacZ)U169 phoA glnV44 Φ80Δ(lacZ)M15 gyrA96 recA1 relA1 endA1 thi-1 hsdR17*). The *TEM-1* gene was under the control of the IPTG-inducible *tac* promoter on pSkunk1, a minor variant of plasmid pSkunk3 (AddGene plasmid #61531) (1). The growth media was LB media supplemented with glucose (2% w/v) and spectinomycin (50 µg/ml) except as noted. Experiments used 100 ml media in 500 ml baffled flasks incubated at 37°C with shaking at 250 rpm. The optical density (OD) of cultures was measured at 600 nm.

**Library construction.** Libraries of all possible single-codon substitutions in TEM-1 were constructed using inverse PCR with mutagenic oligonucleotides obtained from IDT. The oligonucleotides were designed to contain a degenerative codon NNN (hand-mixed) targeted to each codon in the β-lactamase gene. Inverse PCR was performed using the mutagenic oligonucleotides and Phusion™ High-Fidelity PCR MasterMix with HF Buffer (New England Biolabs) following the manufacturer's recommended thermocycling conditions. Amplification of PCR products was verified for each codon site on a 1% TAE gel containing ethidium bromide. Bands were visualized under UV light and imaged with Carestream Gel Logic 112. Inverse PCR was repeated for any sites which did not have a prominent band at ~4 kb as identified by UV transillumination. Successful PCR reactions were pooled together and run in a 1% TAE gel. The desired band was excised from the gel and the linearized DNA extracted using a PureLink™ Quick Gel Extraction Kit (Invitrogen). Plasmid DNA was cleaned using a DNA Clean & Concentrator Kit (Zymo) then phosphorylated using T4 Polynucleotide Kinase (NEB). Phosphorylated DNA was cleaned and concentrated once more before ligation using a T4 DNA Ligase (NEB). The ligation products were transformed into NEB 5-alpha LacI$^q$ competent cells (NEB) and plated on agar dishes made with 200 ml of LB-agar containing 2% w/v glucose and 50 µg/ml spectinomycin. We constructed the library in three separate regions due to read length constraints of Illumina MiSeq. For each region, we estimated that a target of 50,000 transformants would ensure high coverage of the library (2). The transformation step was repeated until an excess of 50,000 transformants had been obtained in total. Random library members were selected and sequenced to verify the presence of single-codon mutations. Colonies were collected from the plates in 15 ml of media containing 15% glycerol. Cells were spun down and the pellet resuspended in a small volume of the supernatant to be stored at -80°C.

In replica 1 of the growth competition experiment, we noticed underrepresented mutations at particular positions within regions 1 and 2 of the library. The aforementioned method was repeated in order to create a supplemental library containing mutations at M3, I5, A17, E28, T29, G41, S53, L57, R65, C77, E89, R94, I95, H96, S98, P107, V108, T109, T181, A184, and R191. This mini-library was spiked into the original library in the replica 2 growth competition experiment.

**Growth competition experiment of the libraries.** For replicas 1 and 2 of region 1 and replica 1 of region 2, frozen library stocks and wildtype cells were diluted to an OD of 0.020 in 100 ml media and incubated at 37°C with shaking until the OD was about 0.5. The wildtype and library cultures were then mixed at a ratio of 5:95 based on OD. Ten ml of this mixture was place on ice and centrifuged at 4000 x g for 10 minutes at 4°C. Plasmid was extracted from pelleted cells using the Plasmid Miniprep Kit (Qiagen) and stored as the initial time point sample. The remaining culture was diluted in a shake flask in media prewarmed to 37°C to obtain a 100 ml culture with an OD of 0.02. At time zero, IPTG was added to a concentration of 1 mM and the flasks were incubated at 37°C with shaking until the OD was approximately 0.640 (10 generations of growth). The culture was then diluted back to an OD of 0.020 in 100 ml of prewarmed media in a separate shake flask and incubated at 37°C with shaking until the OD was approximately 0.640. Ten ml of the culture was placed on ice and then centrifuged at 4000 x g for 10 minutes at 4°C. Plasmid was extracted as the generation 10 sample.

For replica 2 of region 2 and replicas 1 and 2 of region 3, a slightly different protocol was followed that allowed the cells longer time to recover from being frozen before the start of the growth competition. In particular, we suspected that the frozen stocks of the wildtype cells, which had been frozen much longer than the libraries, took more time to recover from being frozen and reach maximal growth rate – more time than the 4-5 generations provided (see analysis of fitness data section). For this modified protocol, frozen library stocks and wildtype cells were diluted into 100 ml media containing 50 µg/ml spectinomycin and 2% w/v glucose and incubated overnight at 37°C. The wildtype and library cultures were then mixed at a ratio of 5:95 based on OD, diluted to an OD of 0.020 in 100 ml media, and incubated at 37°C with shaking until the OD was about 0.5. Ten ml of this mixture was placed on ice before the plasmid was extracted as the initial sample. The remaining culture was diluted to an OD of 0.020 in 100 ml of media in a shake flask, which had been prewarmed to 37°C. With the addition of IPTG, the protocol continued as described above.

Plasmid collected during the growth competition experiment was digested with SphI-HF (NEB) and purified using a DNA Clean & Concentrator Kit (Zymo). Custom Illumina adapter sequences ordered from IDT were added to the linearized DNA for PCR amplification. Adapters contained unique barcodes to identify which region and time point the DNA corresponded to. Proper DNA size was verified on a 1% agarose gel with ethidium bromide. Samples were pooled and submitted for Illumina MiSeq (2 x 300 bp reads) at the Transcriptomics and Deep Sequencing Core facility at Johns Hopkins University.

**Deep sequencing analysis.** Illumina MiSeq reads for each of the three regions were inspected for per base sequence quality using FastQC (3). Paired-end reads were merged using PEAR (4) set to a minimum assembly length of 200 base pairs. Illumina adapters and base pairs outside the desired regions were cropped from the merged reads using Trimmomatic (5) (Region 1-HEADCROP:24, CROP:285; Region 2- HEADCROP:20, CROP:285; Region 3-HEADCROP:24, CROP:291). The resulting trimmed reads were input to Enrich2 (6), which counted the variants for use in calculating selection coefficients and variance. Reads containing bases with a quality score below 20, bases marked as N, or mutations at more than one codon were filtered out.

**Fitness calculation for DMS.** We calculated fitness values using the allele counts tabulated by Enrich2 and the fold increase in the number of cells during the experiment. Our approach to calculating fitness from growth competition experiments using deep sequencing is very similar to (and based on) that of Kowalsky et al. (7) and Rubin et al. (6). The difference is our approach calculates a fitness based on growth rate instead of using an enrichment ratio as a proxy for fitness.

The growth rate of the bacteria ($\mu$) from time 0 to $t$ is proportional to the initial total number of cells ($N_o$) and the final number of cells ($N_f$) at time $t$ according to Equation 1.

$$\mu = \frac{1}{t}\ln\frac{N_f}{N_o} \quad (1)$$

We assumed that optical density $O$ of a culture is linearly proportional to the number of cells, thus

$$N_o = k_o O_o \quad (2)$$
$$N_f = k_f O_f \quad (3)$$

We assumed $k_o = k_f$, and since the culture underwent one *d*-fold dilution during growth

$$\frac{N_f}{N_o} = \frac{O_f d}{O_o} \quad (4)$$

The rate of growth of cells expressing an allele *i* can be written in terms of the sequencing counts of that allele ($c_i$) and the total sequencing count ($c_T$)

$$\mu_i = \frac{1}{t}\ln\left(\frac{\frac{c_{if}}{c_{Tf}}(O_f d)}{\frac{c_{io}}{c_{To}}O_o}\right) \quad (5)$$

We defined the enrichment of allele *i* as

$$\varepsilon_i = \frac{c_{if}c_{To}}{c_{io}c_{Tf}} \approx \frac{(c_{if}+0.5)c_{To}}{(c_{io}+0.5)c_{Tf}} \quad (6)$$

We add the 0.5 to all counts of mutant alleles to assist with alleles that have counts that are zero (i.e. so that a fitness value can still be calculated) (6).

By defining the fold increase in the total number of cells (*r*) as

$$r = \frac{O_f d}{O_o} \quad (7)$$

we rewrote Equation 5 for the rate of growth of cells containing allele *i* by combining it with Equations 6 and 7.

$$\mu_i = \frac{1}{t}\ln(r\varepsilon_i) \quad (8)$$

Since fitness of an allele ($w_i$) is simply the ratio of the growth rate of allele *i* to that of wildtype (wt), fitness will not depend on the time.

$$w_i = \frac{\mu_i}{\mu_{wt}} = \frac{\ln(r\varepsilon_i)}{\ln(r\varepsilon_{wt})} \quad (9)$$

**Statistical treatment of DMS fitness measurements.** The uncertainty in the fitness measurement depends on the sequencing counts, which we assume have a binomial distribution. There are four proportions to consider in calculating the standard error (i.e. the frequencies of the mutant and wildtype at the beginning and end of the experiment). For proportions, the variance $\sigma^2$ depends on the proportion *p* and the number of samples from the population by the following relationship

$$\sigma^2 = \frac{p(1-p)}{n} \quad (10)$$

In our case, $p = c_i/c_T = f_i$ (where $f_i$ is the frequency of allele *i*) and $n = c_T$.

Thus, the variance in the final and initial frequencies of an allele are

$$\sigma_{f_{if}}{}^2 = \frac{\frac{c_f}{c_{Tf}}(1-\frac{c_f}{c_{Tf}})}{c_{Tf}} = \frac{f_f(1-f_f)}{c_{Tf}} \quad (11)$$

$$\sigma_{f_{io}}{}^2 = \frac{\frac{c_o}{c_{To}}(1-\frac{c_o}{c_{To}})}{c_{To}} = \frac{f_o(1-f_o)}{c_{To}} \quad (12)$$

We can rewrite this Equation 5 as

$$\mu_i = \frac{1}{t}\left[\ln\left(rf_{if}\right) - \ln(f_{io})\right] \quad (13)$$

And the fitness as

$$w_i = \frac{\mu_i}{\mu_{wt}} = \frac{\left[\ln\left(rf_{if}\right)-\ln(f_{io})\right]}{\left[\ln\left(rf_{wtf}\right)-\ln(f_{wto})\right]} = \frac{\ln(r\varepsilon_i)}{\ln(r\varepsilon_{wt})} \quad (14)$$

And the variance in the natural log of the frequencies (we assumed the error in *r* is negligible because it appears in the numerator and denominator and the primary source of uncertainty is in the counts)

$$\sigma_{\ln\left(rf_{if}\right)}{}^2 = \frac{\sigma_{f_{if}}{}^2}{f_{if}{}^2} = \frac{\left(1-f_{if}\right)}{c_{if}} \quad (15)$$

$$\sigma_{\ln(f_{io})}{}^2 = \frac{\sigma_{f_{io}}{}^2}{f_{io}{}^2} = \frac{(1-f_{io})}{c_{io}} \quad (16)$$

Thus, the variance in the $\ln(r\varepsilon)$ terms are

$$\sigma_{\ln(r\varepsilon_i)}{}^2 = \frac{\left(1-f_{if}\right)}{c_{if}} + \frac{(1-f_{io})}{c_{io}} \quad (17)$$

$$\sigma_{\ln(r\varepsilon_{wt})}{}^2 = \frac{\left(1-f_{wtf}\right)}{c_{wtf}} + \frac{(1-f_{wto})}{c_{wto}} \quad (18)$$

And the variance in the fitness is

$$\sigma_w{}^2 = w_i{}^2\left[\frac{\frac{\left(1-f_{if}\right)}{c_{if}}+\frac{(1-f_{io})}{c_{io}}}{(\ln r\varepsilon_i)^2} + \frac{\frac{\left(1-f_{wtf}\right)}{c_{wtf}}+\frac{(1-f_{wto})}{c_{wto}}}{(\ln r\varepsilon_{iwt})^2}\right] \quad (19)$$

And the confidence interval is

$$\pm z^* \frac{\sigma_w}{\sqrt{1}} \quad (20)$$

Where $z^*$= 2.576 for the 99% confidence interval.

To calculate *P*-values, we wanted to know if the final frequency of counts of allele *i* relative to wildtype is different than the initial frequency of counts of allele *i* relative to wildtype. Inspecting Equation 9, we see there would be a fitness value of 1.0 if

$$\varepsilon_i = \varepsilon_{wt} \quad (21)$$

Since both sides in Equation 21 have the same total counts, all total count values cancel out and Equation 21 becomes

$$\frac{c_{io}}{c_{wt0}} = \frac{c_{if}}{c_{wtf}} \quad (22)$$

We asked if the left proportion is different than the right proportion. We tested against the null hypothesis that these proportions are the same, which is:

$$H_0: \frac{c_{io}}{c_{wt0}} = \frac{c_{if}}{c_{wtf}} \quad (23)$$

We calculated a *Z*-score for this:

$$Z = \frac{\hat{p}_o - \hat{p}_f}{\sqrt{\hat{p}(1-\hat{p})\left(\frac{1}{c_{wto}} + \frac{1}{c_{wtf}}\right)}} \quad (24)$$

$$\hat{p}_o = \frac{c_{io}}{c_{wto}} \quad (25)$$

$$\hat{p}_f = \frac{c_{if}}{c_{wtf}} \quad (26)$$

$$\hat{p} = \frac{c_{io} + c_{wto}}{c_{if} + c_{wtf}} \quad (27)$$

Equation 24 is the pooled version of this equation, which is used for comparing “no difference” instead of comparing a specific difference.

*P*-values were calculated by determining the area under the curve for that *Z*-score for one tail of the normal distribution (using the NORMSDIST(Z) function in Excel) and then multiplying by 2 because we did a 2-tailed test (i.e. the fitness might be higher or lower than 1).

In order to correct for multiple testing (8), we estimated the number of false positives that would be included at P<0.01 and P<0.001 significance for each replicate of DMS. *P*-values were placed in a column vector and sorted numerically from smallest to largest. Each *P*-value was assigned an index (i) in ascending order and the corresponding q-values calculated:

$$q_i = \frac{p_i \cdot (N)}{i} \quad (28)$$

Where N represents the total length of the column vector.

We then identified the index and q-value corresponding to P-values of 0.01 and 0.001. The number of false positives at that level of significance were estimated for each replicate of DMS:

$$Number\ of\ false\ positives = q_i \cdot (i) \quad (29)$$

Correcting for multiple testing, we estimate that our data would contain approximately 55.0 false positives on average at $P<0.01$ significance or an estimated 5.6 false positives on average at $P<0.001$ significance for a single replica. We chose to report the frequency of mutations having fitness effects that met the *P*-value criteria in both replica experiments. The false positive rate for such analysis is negligible given the high rate of positives.

**Choice of wildtype reference**. For replicas 1 and 2 of region 1 and replica 1 of region 2, we noticed that if we calculated a fitness of all the alleles that were synonyms of TEM-1, the values were 2.2%, 1.5%, and 1.9% higher than 1.0, respectively. We were skeptical of this result. We also noticed an inordinate amount of nonsynonymous alleles with fitness values slightly above 1.0, some of which appeared to be beneficial mutations based on *P* values. We were concerned that this result artifactually arose because we had not allowed enough time for the cells to recover from being stored as frozen stocks at -80°C (see growth competition methods above). We speculated that cells with the wildtype allele, which had been stored separately and had been frozen for a much longer time, might require longer to fully recover than the few hours it took for the culture to reach OD 0.5 during the pre-induction growth period.

To test this idea, for replica 2 of region 2 and replicas 1 and 2 of region 3, we allowed all cultures to grow overnight to saturation before starting the pre-induction growth period on the next day (see growth competition methods). This altered protocol resulted in overall wildtype synonym allele fitness values to be closer to 1.0 (values were 0.7% lower, 0.8% lower and 0.3% higher, respectively). For this reason, we used the wildtype synonym data as a reference to calculate fitness values for replicas 1 and 2 of region 1 and replica 1 of region 2. We used the wildtype allele as the reference to calculate the fitness values of replica 2 of region 2 and replicas 1 and 2 of region 3. As a check on this adjustment, we compared the mean and median difference between fitness values of region 2 for replica 1 (uses wildtype synonym reference) and replica 2 (uses wildtype reference). The mean difference in fitness (for replica 1 minus replica 2) was +0.007 and the median difference was +0.010. Thus, even by using the wildtype synonyms as the reference for replica 1, which lowered all fitness values by 1.5%, fitness values for replica 1 were slightly higher than those of replica 2 (i.e. we did not overcorrect the fitness values of replica 1 by using the wildtype synonyms as the reference).

**Comparison of primary and collateral fitness effects of mutations.** We analyzed the sequencing count data of Stiffler et al (9) to determine the mean growth rate-based primary fitness effect of mutations (**Data S2-S5**). Their experiment was similar to ours but differed in a couple of respects. Both experiments were performed with K-12 *E. coli* in LB media at 37°C. Differences from our experiment include that TEM-1 was under its native promoter on pBR322 (instead of a *tac* promoter on a lower copy plasmid), the cells were MegaX DH10B T1 cells (instead of NEB 5-alpha LacI$^q$), the cultures were 1 ml in 96-well plates (instead of 100 ml in 500 ml shake flasks), and the number of cultures doublings was 3-4 (instead of ~10).

For our analysis of their data, fitness values and associated statistics were calculated as was done with collateral fitness effects except that the comparison was between final allele frequency without Amp and final allele frequency with Amp (instead of between initial and final allele frequency of the same culture). The total sequencing reads per Amp condition in their experiments were ~3.5-3.9 million (replica 1) and 8.5-17 million (replica 2). The total sequencing reads per replica in our experiment were 3.8-4.5 million (replica 1) and 3.5-3.7 (replica 2) million. As the total counts are similar for both, a comparison of the relative frequency of primary and collateral fitness effects is justified.

For the comparison of primary and collateral fitness effects we used the weighted mean fitness values and their corresponding 99% confidence interval ($n$=2) to classify a mutation as deleterious, neutral, or cannot be determined. Deleterious mutations were those whose upper bound of the fitness value was <0.98. Neutral mutations were those whose lower bound of the fitness value was >0.98. Mutations were classified as having collateral only effects if their collateral effects were deleterious and the primary effects were neutral. Mutations were classified as having primary only effects if their primary effects were deleterious and the collateral effects were neutral. Mutations had both effects if both primary and collateral effects were deleterious.

**Mutant construction.** A set of 37 mutants was constructed to confirm collateral fitness effects and investigate their mechanisms. We used the QuikChange Lightning Kit (Agilent) to introduce the mutations in α-helices and stop codon mutations. We used inverse PCR to introduce mutations in the signal sequence and mutations involving cysteines. We also used inverse PCR to construct control plasmid pSkunk *Δ*TEM-1 that had the coding region of the TEM-1 gene deleted. We transformed all mutant plasmids into NEB 5-alpha $LacI^q$ chemically competent cells (NEB). Cells were plated on LB-agar plates containing 50 μg/ml spectinomycin and incubated overnight at 37°C. We verified the mutations by Sanger sequencing.

**Monoculture growth assay for fitness.** Cultures were grown exactly the same as in the growth competition experiments for DMS except the cultures were monocultures and not mixtures of wildtype (WT) and mutant alleles. The optical density was measured roughly every 30 minutes for six hours with a dilution at about 3 hours.

We assumed that the correlation between OD and cell number was the same for cells expressing any TEM-1 allele. We calculated the fitness directly from the starting and final OD's ($O_o$, $O_f$) of the mutant and wildtype cultures and the dilution factor $d$.

$$w_i = \frac{\mu_i}{\mu_{wt}} = \frac{\left(\ln\frac{O_f d}{O_o}\right)_i}{\left(\ln\frac{O_f d}{O_o}\right)_{wt}} \quad (30)$$

The initial OD was calculated based on the OD before dilution and the dilution factor, as the error in OD relative to the OD value was large at this low OD value (0.020). The fitness value in Equation 30 is thus based on the mean growth rate over about 10 generations.

Fitness values based on steady-state growth rates were calculated from the OD data as a function of time between about 4 and 6 hours post-induction using OD values less than 0.3, since *E. coli* steady-state growth stops at about an OD of 0.3 in LB media (10). The steady-state growth rate is the slope of the linear fit of ln(OD) vs. time. We calculated fitness values as the ratio of the steady state growth rate of cells expressing allele *i* to the steady state growth rate of cells expressing wildtype TEM-1.

**Co-culture growth assay.** Frozen stocks of wildtype, A134G, and L148R cells were used to inoculate 10 ml of media containing 50 μg/ml spectinomycin and incubated overnight at 37 °C. The overnight cultures were diluted by adding 5 ml culture to 95 ml media in a sterile flask and then incubated at 37°C with shaking until the OD was about 0.5. These cultures were diluted to an OD of 0.02. Co-cultures were made by mixing 50 ml of wildtype monoculture with 50 ml of mutant monoculture and adding IPTG to a final concentration of 1 mM. A 1 ml sample of the resulting culture was collected to be used as described in the subsequent two subsections. Cultures were grown the same as in the growth competition experiment for DMS over the remainder of the experiment with a 1 ml sample of co-culture collected after 10 generations of growth.

**Determination of fitness from the co-culture growth assay using plating in the presence and absence of Amp.** The co-culture samples collected at the initial time point and after 10 generations of growth were diluted to an OD of $7.23 \times 10^{-6}$ immediately after collection. Samples were plated by adding 100 µl of culture to agar plates made with 20 ml of LB-agar, 50 µg/ml spectinomycin, 2% w/v glucose, and 1 mM IPTG then spreading evenly with glass beads. Six plates were made for each co-culture at the initial time point and twenty plates were made for each co-culture at generation 10. Half of each set of plates contained 2048 µg/ml ampicillin. Due to the large deleterious effect on Amp resistance for the A134G and L148R mutations, only wildtype cells could grow on plates containing this concentration of Amp. Plates were incubated overnight at 37°C. Colonies were counted using a colony counter (Bel-Art). We calculated the fitness using the starting and final ODs and the average colony forming units (CFU) on plates with Amp ($CFU_{+Amp}$) and without Amp ($CFU_{-Amp}$) for each set of plates.

$$O'_{f,wt} = O_f d \frac{CFU_{+Amp}}{CFU_{-Amp}} \quad (31)$$

$$O'_{f,i} = O_f d \frac{CFU_{-Amp} - CFU_{+Amp}}{CFU_{-Amp}} \quad (32)$$

$$w_i = \frac{\ln\left(\frac{O'_{f,i}}{O'_{o,i}}\right)}{\ln\left(\frac{O'_{f,wt}}{O'_{o,wt}}\right)} \quad (33)$$

The values for $O'_{o,i}$ and $O'_{o,wt}$ are both 0.01 since the co-culture contained an even proportion of wildtype and mutant at its initial OD of 0.02.

**Determination of fitness from the co-culture growth assay using Sanger DNA sequencing.** Co-cultures were grown as described in the co-culture growth assay. The samples were centrifuged and plasmid was isolated from the pelleted cells using the Plasmid Miniprep Kit (Qiagen) for Sanger sequencing. The resulting chromatogram files were analyzed with Sequence Scanner Software v2 to integrate the area under the curve of the wildtype base pair's peak and the mutant's peak at the site of the single base-pair mutation. A fraction of total peak area for WT and mutant nucleotides were calculated. These fractions were compared to a standard curve to determine the fraction of WT ($F_{wt}$) and mutant ($F_i$) cells at the beginning and end of the growth competition. The standard curve was constructed by sequencing plasmid DNA isolated from mixtures of uninduced WT and mutant cells at set ratios ranging from 50% wildtype to 90% wildtype by 5% intervals (ratios based on OD). We calculated the fitness by substituting the values calculated in Equations 34-35 in Equation 33.

$$O'_{wt} = OdF_{wt} \quad (34)$$

$$O'_i = OdF_i \quad (35)$$

**Cell fractionation and analysis by SDS-PAGE and western blot.** We followed the BugBuster reagent protocol for cell fractionation (Millipore Sigma) except as noted below. Cell cultures were grown as described in the monoculture growth assays. At the end of the growth period, two 40 ml samples of culture were collected in Nalgene centrifuge bottles and centrifuged at 3000 x g and 4°C for 20 minutes. The supernatant was discarded. One of the bottles from each cell culture was frozen overnight at -80°C. In the other bottle, 200 µl of TSE buffer was added and gently swirled around until the pellet was resuspended. The cells were then placed on ice for 30 minutes before the cell suspension was transferred to a microcentrifuge tube and centrifuged at 16,000 x g for 30 minutes at 4°C. Approximately 190 µl of supernatant was removed as the periplasmic fraction. The next day, the Nalgene bottle stored at -80°C was thawed and the mass of the pellet recorded. The pellet was then resuspended in room temperature BugBuster reagent containing 1% v/v protease inhibitor (Thermofisher) and 0.1% v/v Benzonase Nuclease (Millipore Sigma). For each gram of cell pellet, 5 ml of BugBuster master mix was added. The cell suspension was transferred to a microcentrifuge tube and then incubated on a shaking platform for 20 minutes at room temperature before being centrifuged at 16,000 x g for 20 minutes at 4°C. The supernatant was recovered and stored as the soluble fraction, and the remaining pellet was stored on ice. To each ml of BugBuster reagent, 1 µl of 1 KU/ µl rLysozyme (Novagen) was added. The pellet on ice was resuspended in 200 µl of BugBuster with rLysozyme and vortexed. After a five-minute incubation at room temperature, 200 µl of BugBuster diluted tenfold in deionized water was added to the suspension and vortexed for one minute. The suspension was centrifuged at 16,000 x g for 15 minutes and 4°C and the supernatant saved as the membrane fraction. The remaining pellet was washed with 100 µl of diluted BugBuster reagent, vortexed, and centrifuged again. The supernatant was added to the membrane fraction, whereas the pellet was resuspended in 50 µl of diluted BugBuster and stored as the inclusion body fraction.

Concentrations of the protein fractions were measured using the DC Assay Kit (Bio-Rad). A standard curve was formed with BSA ranging from 0.25 mg/ml to 2 mg/ml.

Reducing gel samples were prepared with 15 µg of protein, 4X LDS sample buffer (Invitrogen), 10X sample reducing agent (Invitrogen), and the remainder of the maximum well loading volume as nuclease free water. Samples were spun down in a microcentrifuge and placed in a 95°C heat block for 5-10 minutes. Following denaturation, the samples were cooled on ice for three minutes then allowed to cool at room temperature for 15 minutes before loading. Non-reducing samples were prepared in the same manner but with the reducing agent omitted.

Fractions were analyzed by SDS-PAGE gel electrophoresis using 4-12% NuPage Bis-Tris gels (Thermo Fisher). Gels were run in MOPS running buffer at 200 V. Protein was transferred to a PVDF membrane using a semi-dry electrophoretic transfer cell set at 15 V for 15 minutes. Gels were stained with Coomassie Brilliant Blue R-250 (Bio-Rad) and then destained with destaining solution (50% methanol, 40% water, 10% glacial acetic acid) following transfer in order to verify even loading. PVDF membranes were placed in a solution of 5% nonfat dry milk (Bio-Rad) in TBS-T for one hour, incubated overnight in rabbit Anti-Beta Lactamase polyclonal antibody (AB3738-I; Millipore Sigma), and incubated in goat anti-rabbit IgG (H+L)-HRP conjugate (Bio-Rad) for 90 minutes. Imaging of the membrane was done by western blot using a Clarity Western ECL Substrate Kit (Bio-Rad) and captured on a Bio-Rad ChemiDoc Molecular Imager.

**Nitrocefin assay for TEM-1.** Whole-cell lysate collected during cell fractionation was measured for total protein concentration by DC Protein Assay (Bio-Rad). A volume corresponding to 250 µg of total soluble protein was loaded in a quartz cuvette with 200 µl of 500 mM phosphate buffer (pH 7.4) and deionized water to a final volume of 1,980 µl. The cuvette was placed in a Cary 50 UV-Vis Spectrophotometer with a micro stir bar. A high stir rate and temperature of 25°C were maintained using a Single Cell Peltier Accessory (Agilent). Absorption data at 486 nm was collected for one minute after adding 20 µl of 10 mM nitrocefin (TOKU-E) to the cuvette. The differences in slope within the initial linear range of absorption data was used to calculate the relative levels of TEM-1 in the whole-cell lysates.

**RNA-Seq Sample Preparation and Analysis.** Cell cultures were prepared following the monoculture growth assay protocol. Controls consisted of cells with the wildtype allele that were not induced with IPTG as well as cells harboring pSkunkΔTEM-1 that were induced with IPTG. Growth was halted at about 330 minutes (i.e. at or before an OD of 0.3) to ensure the cells were still in exponential growth phase based on OD measurements. Microcentrifuge tubes that had been chilled on ice were filled with 1.5 ml of bacteria culture and centrifuged at 6000 x g and 4°C for 5 minutes. Supernatant was decanted and the cell pellet resuspended in 200 µl of Max Bacterial Enhancement Reagent (Invitrogen) that had been preheated to 95°C. The tubes were incubated at 95°C for 4 minutes before 1 ml of TRIzol Reagent (Invitrogen) was added. Sample lysis, separation of the RNA aqueous phase, and subsequent precipitation and wash steps were performed according to the TRIzol user manual. The resulting RNA pellet was resuspended in 50 µl of nuclease-free water. A second RNA precipitation was performed to remove residual salt as recommended by the MICROBExpress Bacterial mRNA Enrichment Kit (Invitrogen). RNA was resuspended in 0.1X TE buffer (NEB). Concentration of RNA was measured by NanoDrop spectrophotometer. RNA integrity was examined by native agarose gel electrophoresis with 300 ng of total RNA loaded in each lane of a 1.0% TAE gel containing ethidium bromide. mRNA enrichment was completed using the MICROBExpress Bacterial mRNA Enrichment Kit. The enriched mRNA was treated with DNase using the TURBO DNA-free Kit (Invitrogen). Double-stranded cDNA was synthesized, purified, and end prepped with the NEBNext Ultra II Directional RNA Library Prep Kit with Sample Purification Beads (NEB). NEBNext Adaptor for Illumina (NEB) was ligated to the cDNA. The ligation reactions were purified using NEBNext Sample Purification Beads and PCR enriched using a unique index primer from the NEBNext Multiplex Oligos for Illumina Kit (NEB) for each sample. PCR products were cleaned with NEBNext Sample Purification Beads and pooled for Illumina Tru-Seq (2 x 75 bp reads) at the Transcriptomics and Deep Sequencing core at Johns Hopkins University.

Reads were analyzed for per base sequence quality using FastQC (3). We indexed the genome of NEB 5-alpha cells (NCBI Reference Sequence: NZ_CP017100.1) and mapped our paired end reads to the transcriptome using STAR (11). For quantifying and tabulating transcript abundance, we used featureCounts (12). Normalized counts per million for the two biological replicates of RNA-Seq were tabulated with edgeR (13). RNA-Seq sequencing counts, fold-difference in gene expression, and associated statistics are provided as **Data S6.**

**Calculation of fold-difference in gene expression.** Values of fold-difference in gene expression ($g_i$) were calculated using the counts for an individual gene ($c_i$) and total sequencing counts ($c_T$).

$$g_i = \frac{c_i c_{T,wt}}{c_{wt} c_{T,i}} \approx \frac{(c_i+0.5)c_{T,wt}}{(c_{wt}+0.5)c_{T,i}} \quad (36)$$

**Statistical treatment of fold-difference in gene expression.** The uncertainty in fold-difference depends on the proportion of total sequencing counts that correspond to an individual gene in both the wildtype and mutant-containing cells. For binomial distributions, a proportion *p* has a variance $\sigma^2$ of

$$\sigma^2 = \frac{p(1-p)}{n} \quad (37)$$

In our case, $p = c_i / c_{i,T} = f_i$ (where $f_i$ is the frequency of counts corresponding to a particular gene in cells with mutation *i*) and $n = c_{i,T}$.

Thus, the variance arising from the mutant and wildtype sequencing counts would be

$$\sigma_{f_{wt}}{}^2 \approx \frac{\frac{c_{wt}+0.5}{c_{wt,T}}\left(1-\frac{c_{wt}+0.5}{c_{wt,T}}\right)}{c_{wt,T}} = \frac{f_{wt}(1-f_{wt})}{c_{wt,T}} \quad (38)$$

$$\sigma_{f_i}{}^2 \approx \frac{\frac{c_i+0.5}{c_{i,T}}\left(1-\frac{c_i+0.5}{c_{i,T}}\right)}{c_{i,T}} = \frac{f_i(1-f_i)}{c_{i,T}} \quad (39)$$

And the variance in the fold-difference is

$$\sigma_{g_i}{}^2 = g_i{}^2\left(\frac{\sigma_{f_{wt}}{}^2}{f_{wt}{}^2} + \frac{\sigma_{f_i}{}^2}{f_i{}^2}\right) = g_i{}^2\left(\frac{1}{c_{wt}+0.5} - \frac{1}{c_{wt,T}} + \frac{1}{c_i+0.5} - \frac{1}{c_{i,T}}\right) \quad (40)$$

The *Z*-score was calculated using equations analogous to those used in analysis of DMS data.

$$Z = \frac{\hat{p}_i - \hat{p}_{wt}}{\sqrt{\hat{p}(1-\hat{p})\left(\frac{1}{c_{i,T}}+\frac{1}{c_{wt,T}}\right)}} \quad (41)$$

$$\hat{p}_i = \frac{c_i}{c_{i,T}} \quad (42)$$

$$\hat{p}_{wt} = \frac{c_{wt}}{c_{wt,T}} \quad (43)$$

$$\hat{p} = \frac{c_i + c_{wt}}{c_{i,T}+c_{wt,T}} \quad (44)$$

*P*-values were calculated using the NORMSDIST(Z) function in Excel as described for the DMS fitness calculations.

Genes and pathways of interest were identified from the set of values in which fold-difference in gene expression differed by >2-fold and *P*<0.001. We then created an ordered list of these fold-difference values and searched for patterns across interconnected genes using EcoCyc (14) and previous studies of stress pathways.

**Colony dye-uptake experiments.** LB media containing 50 μg/ml spectinomycin and 2% w/v glucose was inoculated from frozen stocks of wildtype TEM-1 or mutant-containing cells and incubated overnight at 37°C with shaking. Cultures were diluted to an OD of approximately 7.23x10$^{-6}$ and plated on 20 ml of agar containing 1 mM IPTG, 2% w/v glucose, 50 μg/ml spectinomycin, and 150 μg/ml Congo Red or 25 μg/ml Toluidine Blue-O. The plates were incubated at 37°C for 48 hours and imaged at 24 and 48 hours.

**Data deposition.** DMS sequencing data is available at BioProject (www.ncbi.nlm.nih.gov/bioproject) using accession number PRJNA565437. RNA-Seq sequencing data is available at Gene Expression Omnibus at NCBI (www.ncbi.nlm.nih.gov/geo/) using accession number GSE137362. The processed data are also provided in Excel files as **Data S1**, and **Data S6**.

**Supplementary Text**

Additional changes in gene expression

The other main class of differentially expressed genes were operons regulated by FNR, which regulates expression of a number of genes involved in anaerobic respiratory pathways (15), including genes upregulated when *E. coli* transitions from aerobic to micro-aerobic conditions in the presence of nitrate (16) (**Fig. S19**). We believe these gene expression changes are unrelated to the mutations. Both replica cultures expressing unmutated TEM-1 had elevated expression of these genes. However, cultures expressing the TEM-1 mutants and negative controls show variability in whether these genes' expression were elevated, even between replica experiments. We suspect this reflects flask-to-flask variability in oxygen availability to the culture late in the exponential phase or during sample processing before cell lysis. These differences in gene expression did not correlate with changes in the stress response pathways. These genes were differentially expressed among replicas of A11D cells and among replicas of control cells lacking TEM-1 expression (**Fig. S19**), yet these cells had no differences in the expression of stress response pathways. We regard the differential expression of these genes as an experimental artifact unrelated to the mutations' effects on gene expression.

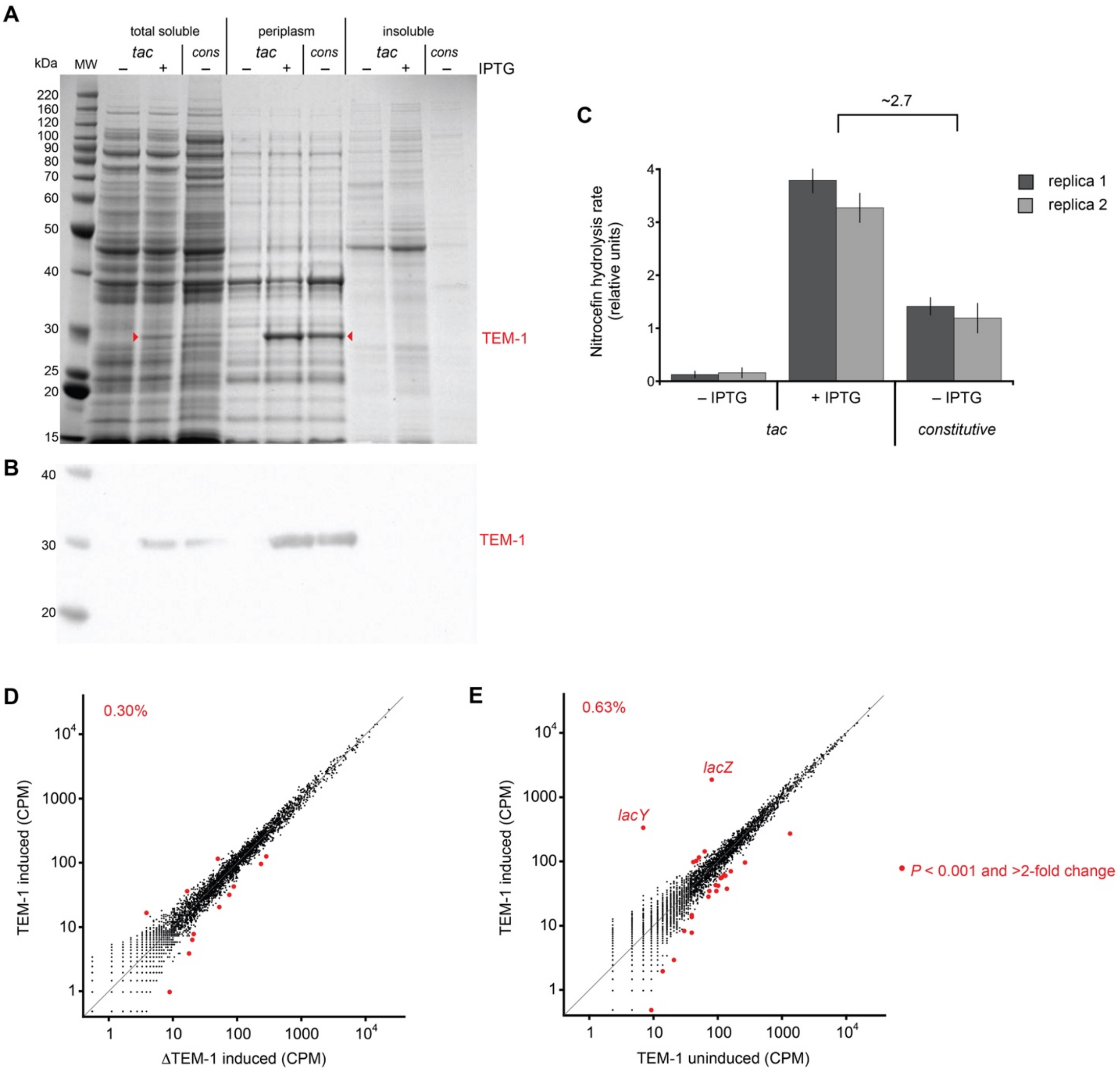


**Figure S1. Production of TEM-1 and its effect.** (**A**) SDS-PAGE gel and (**B**) western blot of cell extracts from NEB 5-alpha F'lacI$^q$ *E. coli* cells expressing TEM-1 from the *tac* promoter on pSkunk1 or the constitutive *TEM-1* promoter (*cons*) of pBR322. One mM IPTG induced expression from the *tac* promoter. The expected size of mature TEM-1 is 28.9 kDa. (**C**) Quantification of TEM-1 protein levels in the total soluble fraction of cell lysates using the nitrocefin colorimetric assay for β-lactamase activity. Initial rates of 100 μM nitrocefin hydrolysis were measured using equal concentrations of total soluble protein. The error bars in the two biological replica experiments represent the standard deviation of the assay (*n*=3). RNA-Seq results showing the effect of TEM-1 on gene expression using (**D**) a plasmid with the *TEM-1* gene deleted as the comparison and (**E**) a culture lacking the inducer IPTG as the comparison. IPTG induces the *lacZYA* operon. The *lacA* gene was differentially expressed in (E) but is not shown because it had zero CPM in the uninduced sample (it had 73 CPM in the induced sample). Data indicated with a larger, red circle indicates genes with at least a 2-fold change in expression and $P < 0.001$ (*Z*-test). The percentage of genes that meet these criteria is indicated on each plot. CPM = counts per million.

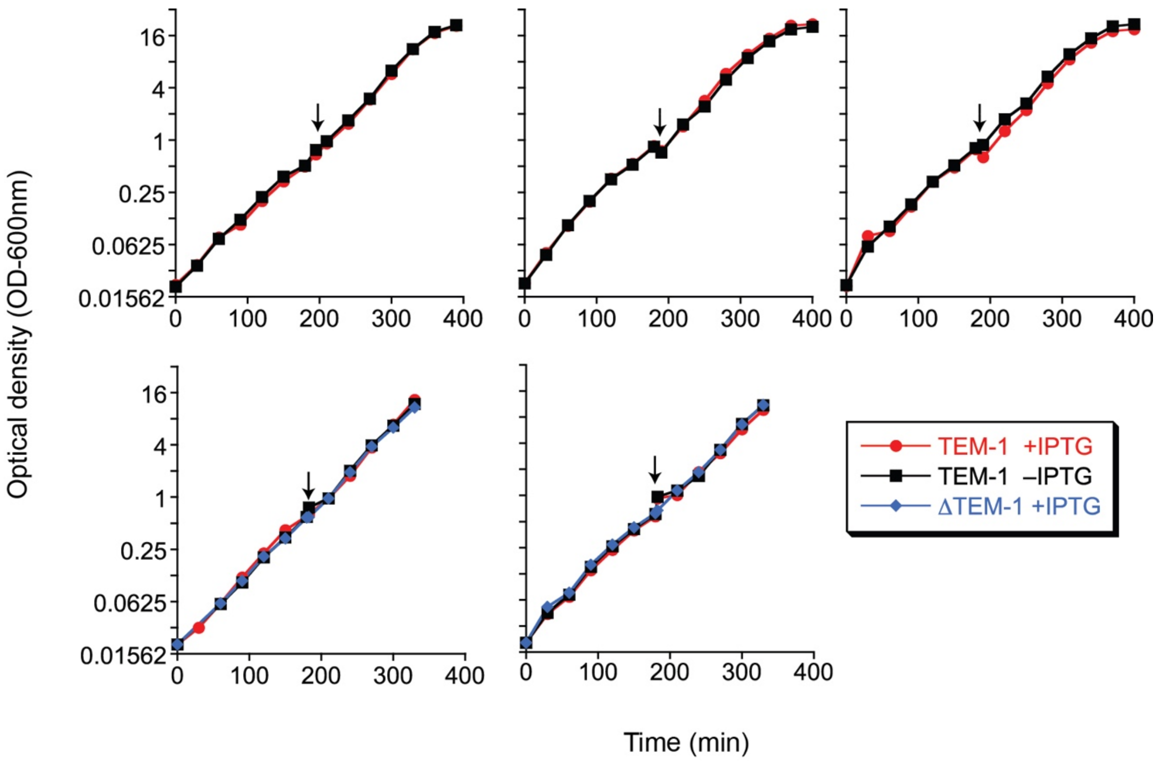


**Figure S2. Effect of TEM-1 expression on fitness.** Five replica growth experiments are shown. At zero minutes, exponentially growing cultures in LB media lacking IPTG were diluted to an optical density of 0.020 and then divided into two separate flasks. IPTG (1 mM final concentration) was added to one flask and the growth was monitored by optical density at 600 nm (OD-600nm) for about 6 hours. At about 3 hours (arrow), the culture was diluted into fresh media (with or without IPTG as appropriate). OD measurements after this dilution are relative values in which the actual OD value is multiplied by the fold-dilution. In the two replica experiments in the second row, an IPTG-containing culture of cells with a plasmid in which the TEM-1 gene had been deleted (ΔTEM-1) was included as a comparison. Based on the OD at the end of the experiment and the number of generations, the fitness effects of inducing TEM-1 expression in the five experiments were -0.5%, +1.5%, -1.9%, +1.7%, and -1.9%. The mean value was -0.2 ± 1.8% (n=5). The differences might partially reflect the precision of the dilution at about 3 hours (i.e. differences on the order of 1% in the accuracy of the dilution between the two flasks). We concluded that the expression of TEM-1 has little, if any, fitness effect relative to the magnitude of most of the collateral fitness effects studied in this work.

**A**

| Mutation | Mutant Codon | Fitness (DSM) replica 1 | Fitness (DSM) replica 2 | Fitness (mean growth rate) |
|---|---|---|---|---|
| F8S | AGC | 1.03 ± 0.05 | 0.94 ± 0.07 | |
| | AGT | 1.00 ± 0.04 | 1.00 ± 0.06 | |
| | TCA | 1.00 ± 0.04 | 1.01 ± 0.05 | |
| | TCC | 0.99 ± 0.04 | 0.96 ± 0.07 | |
| | **TCG** | **0.71 ± 0.03** | **0.71 ± 0.04** | **1.00** |
| | TCT | 1.00 ± 0.11 | 1.08 ± 0.14 | |
| A11T | **ACA** | **0.70 ± 0.03** | **0.73 ± 0.03** | **1.00** |
| | ACC | 1.03 ± 0.03 | 0.99 ± 0.04 | |
| | ACG | 1.00 ± 0.03 | 0.98 ± 0.04 | |
| | ACT | 1.03 ± 0.02 | 1.00 ± 0.04 | |
| F16R | AGA | 1.00 ± 0.05 | 1.01 ± 0.05 | |
| | **AGG** | **0.85 ± 0.02** | **0.81 ± 0.02** | **0.98** |
| | CGA | 1.02 ± 0.04 | 1.03 ± 0.05 | |
| | CGC | 1.01 ± 0.03 | 1.02 ± 0.04 | |
| | CGG | 1.01 ± 0.03 | 0.99 ± 0.04 | |
| | CGT | 1.03 ± 0.03 | 1.04 ± 0.05 | |
| Q39R | **AGA** | **0.78 ± 0.02** | **0.79 ± 0.03** | **0.98** |
| | AGG | 0.99 ± 0.02 | 0.96 ± 0.03 | |
| | CGA | 1.02 ± 0.03 | 1.02 ± 0.04 | |
| | CGC | 1.00 ± 0.03 | 1.04 ± 0.04 | |
| | CGG | 1.01 ± 0.03 | 1.00 ± 0.03 | |
| | CGT | 1.02 ± 0.03 | 1.02 ± 0.04 | |
| V80R | AGA | 0.95 ± 0.05 | 0.84 ± 0.09 | |
| | **AGG** | **0.67 ± 0.07** | **0.67 ± 0.11** | **0.98** |
| | CGA | 0.96 ± 0.09 | 0.96 ± 0.12 | |
| | CGC | 0.94 ± 0.16 | 1.11 ± 0.35 | |
| | CGG | | | |
| | CGT | 0.95 ± 0.12 | 0.76 ± 0.23 | |
| L81R | **AGA** | **0.74 ± 0.03** | **0.75 ± 0.04** | **0.98** |
| | AGG | 0.90 ± 0.04 | 0.95 ± 0.06 | |
| | CGA | 0.99 ± 0.07 | 0.95 ± 0.10 | |
| | CGC | 0.92 ± 0.09 | 0.84 ± 0.12 | |
| | CGG | 0.96 ± 0.05 | 0.84 ± 0.09 | |
| | CGT | 0.62 ± 0.13 | 0.36 ± 0.53 | |
| S124G | GGA | 0.95 ± 0.04 | 1.00 ± 0.04 | |
| | **GGC** | **0.50 ± 0.04** | **0.52 ± 0.03** | |
| | GGG | 1.02 ± 0.02 | 1.00 ± 0.03 | |
| | GGT | 0.99 ±0.03 | 1.02 ± 0.03 | 1.01 ± 0.01 |

**B**

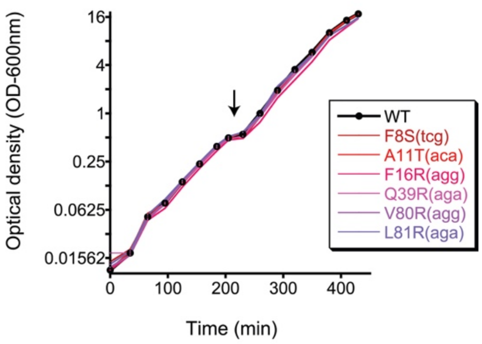


**Figure S3. Effect of synonymous codons on fitness.** (**A**) List of amino acid substitutions that are examples of the rare cases in which one particular codon (noted in red) had a significantly deleterious fitness effect (as measured in replica DMS experiments), but the remaining codons did not. The uncertainties are 99% confidence intervals. These putative deleterious mutations were tested in monoculture growth assays and found to have negligible effects on fitness. The growth assays were performed once except for S124G(ggt) (for which the error is the standard deviation, *n*=3). (**B**) Data from the single replica monoculture growth assays. Growth conditions were the same as in the DMS experiments. At zero minutes, exponentially growing cultures of cells containing the indicated alleles were diluted in LB media to an optical density of 0.020 and IPTG (1 mM final concentration) was added. Growth was monitored by optical density (OD) measured at 600 nm for about 6 hours. At about 3 hours (arrow), the culture was diluted into fresh media (with or without IPTG as appropriate) when the wildtype culture had reached an OD-600nm of approximately 0.64. OD measurements after this dilution are relative values in which the actual OD value is multiplied by the fold-dilution. The fitness values for the growth assay are based on the OD at the beginning and end of the experiment and the number of generations.

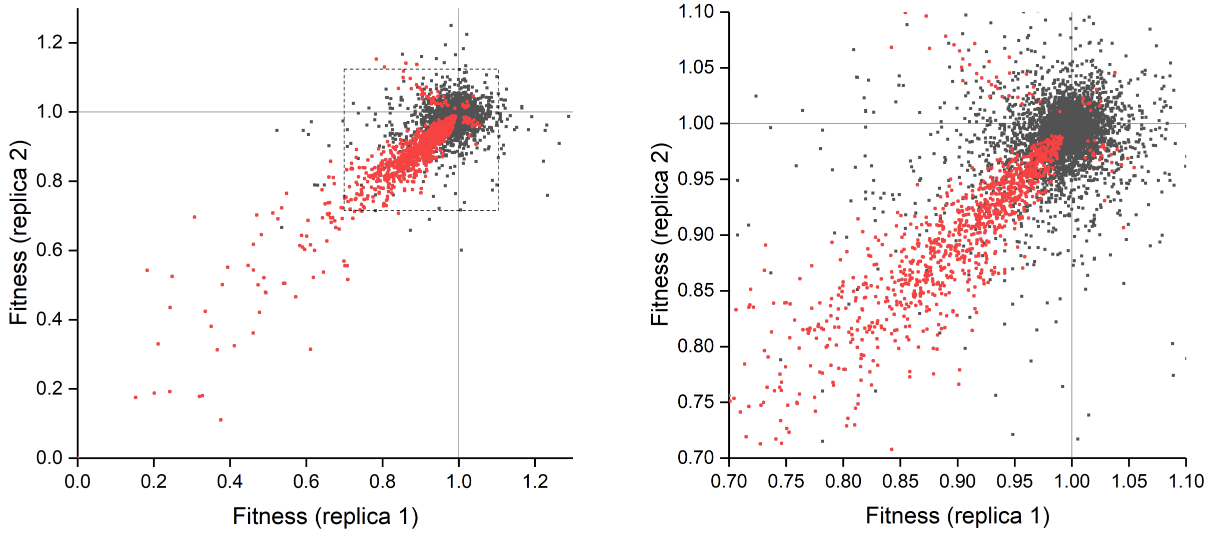


**Figure S4. Correspondence in fitness values between biological replica of DMS experiments.** Deleterious fitness values in replica 1 that were significantly less than 1.0 tended to have fitness values less than 1.0 in the second replica experiment. The right panel is a detail of the left in the region outlined by the dashed box. Data indicated in red indicates fitness measurements that were different than 1.0 at P<0.01 in both replicas.

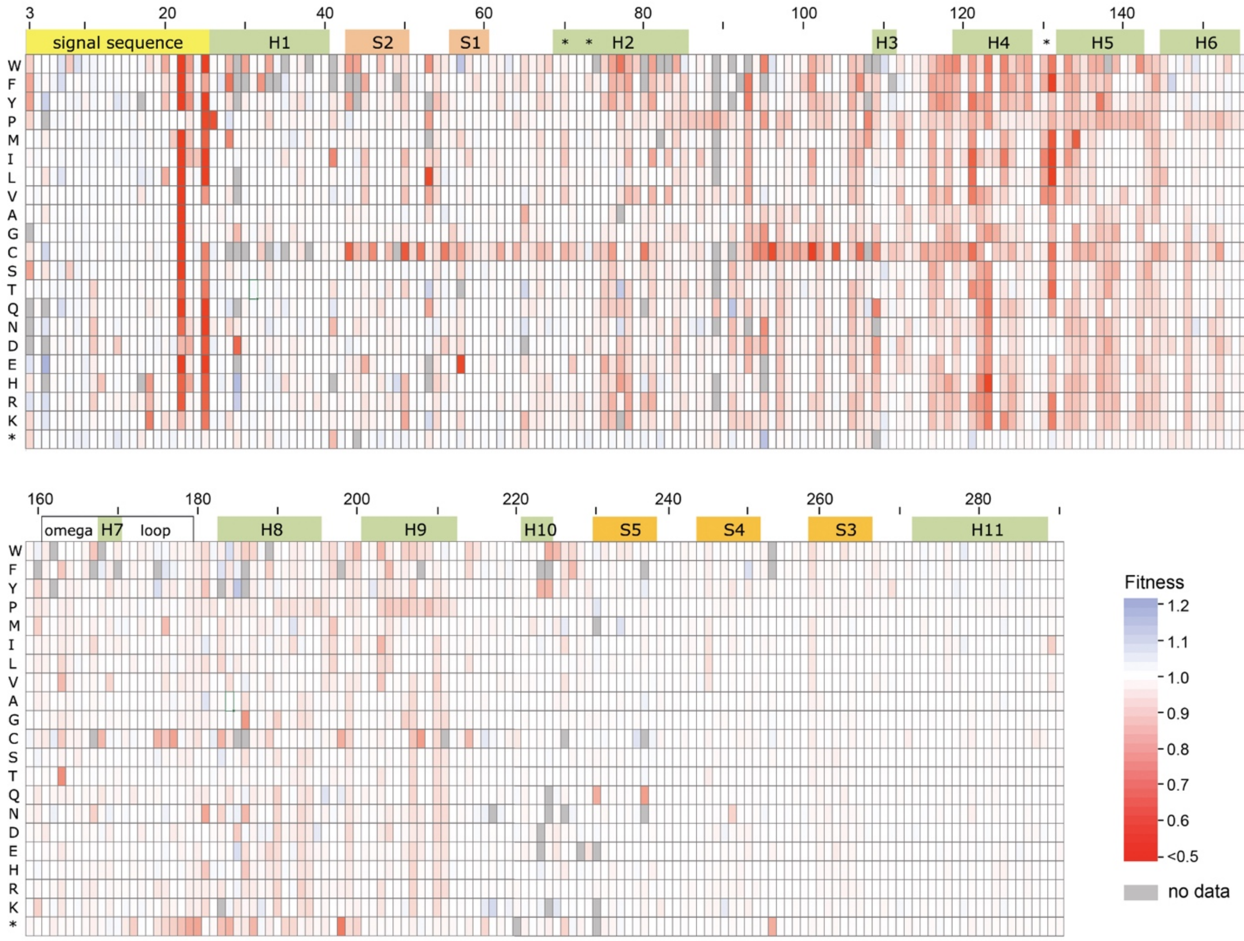


**Figure S5. Weighted mean fitness measurements.** Heat map of weighted mean fitness values for cells expressing TEM-1 with the indicated mutations. The data is the weighted mean of two replica experiments conducted at 37°C in LB media supplemented with 2.0% glucose and 1 mM IPTG. Mutations with no fitness measurement in either replicate are shown in grey. The Ambler numbering system (17) for class A $\beta$-lactamases is used. Regions corresponding to the signal sequence (yellow), $\alpha$-helices (green), $\beta$-strands (orange), the $\Omega$-loop (white) and three key active site residues (*) are shown. Tabulated fitness values are provided as **Supplementary Data S1**.

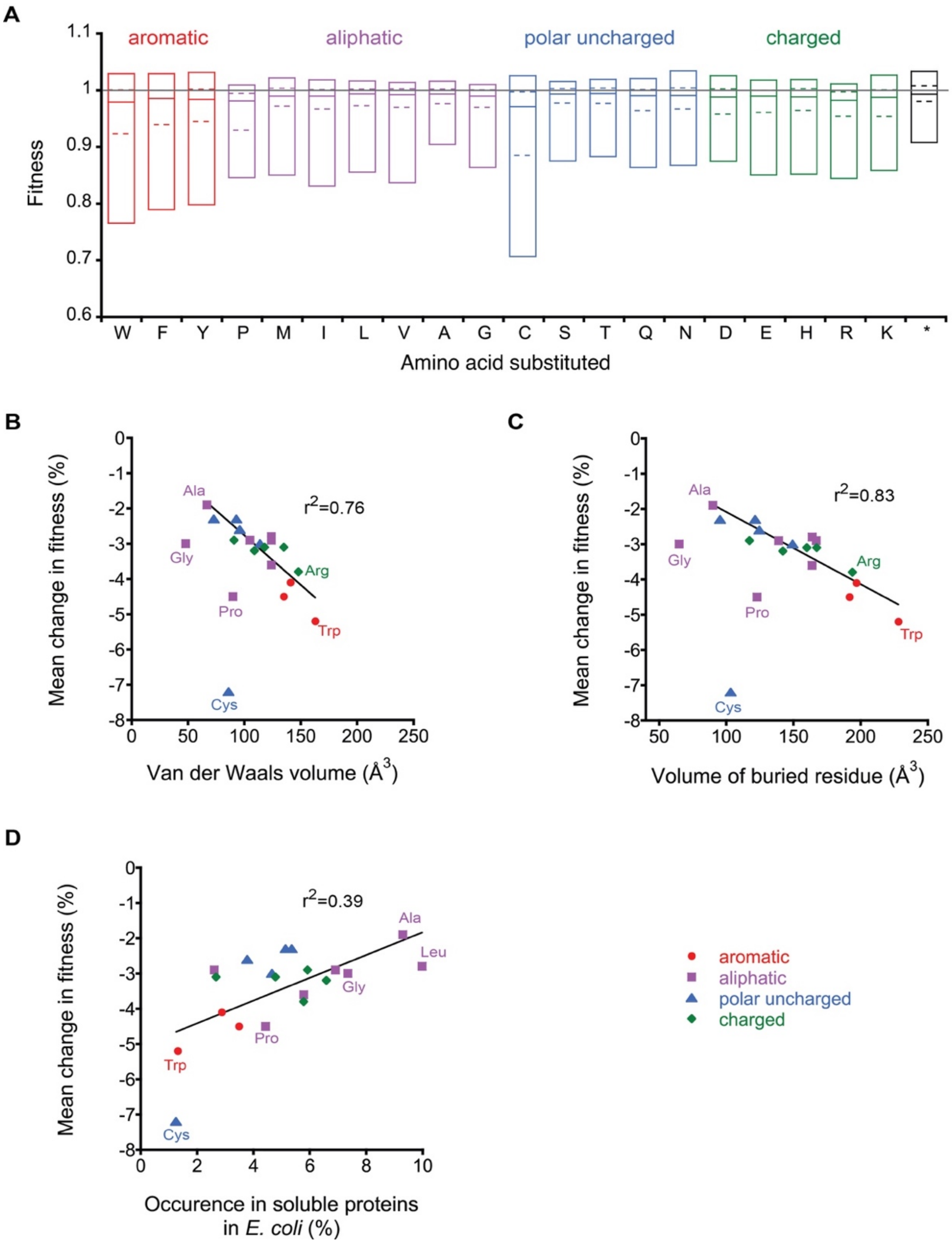


**Figure S6. Fitness effects by mutant amino acid.** (**A**) Fitness values by the amino acid substituted. Box blots indicate the median (solid line), the 25th and 75th percentile (dashed line), and the 5$^{th}$ and 95$^{th}$ percentile (the box). Correlation between the mean change in fitness upon substituting the amino acid in TEM-1 and (**B**) the Van der Waals volume of the amino acid, (**C**) the average volume of the buried residue (18), and (**D**) the natural occurrence of amino acids in soluble proteins of *E. coli* (19). For (B) and (C), the linear fits exclude Gly, Pro, and Cys. The rationale for the exclusion is that these residues have unique attributes that could impact fitness (i.e. cysteine's reactivity, proline's constraint of the peptide backbone, and glycine's lack of a side chain providing peptide backbone flexibility).

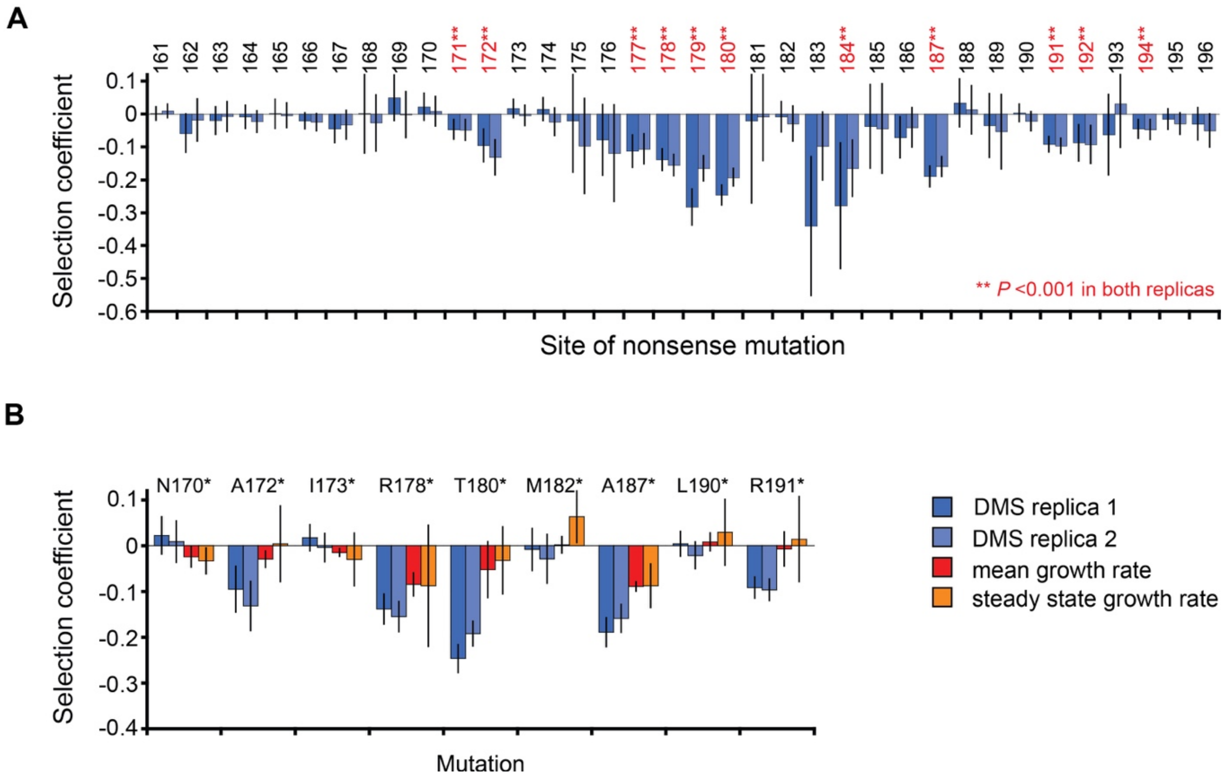


**Figure S7. Collateral fitness effects of nonsense mutations in the region 161-196.** (**A**) Pattern of fitness effects caused by nonsense mutations as measured by two replica DMS growth competition experiments. Error bars are 99% confidence intervals. Positions with fitness effects are labeled in red (**, *P*<0.001 in both experiments). (**B**) Fitness effects caused by select mutations are presented as the percent change in fitness caused by the mutation as measured by DMS (blue, two biological replicates), mean growth rate over 6 hours post-induction (red; $n \geq 3$), and steady state growth rate between 4 and 6 hours post-induction ($n \geq 3$). Error bars represent 99% confidence intervals. For the mean and steady state growth rates, cells expressing the wildtype and mutant proteins were grown in separate flasks for ten generations of wildtype growth as monitored by OD-600nm.

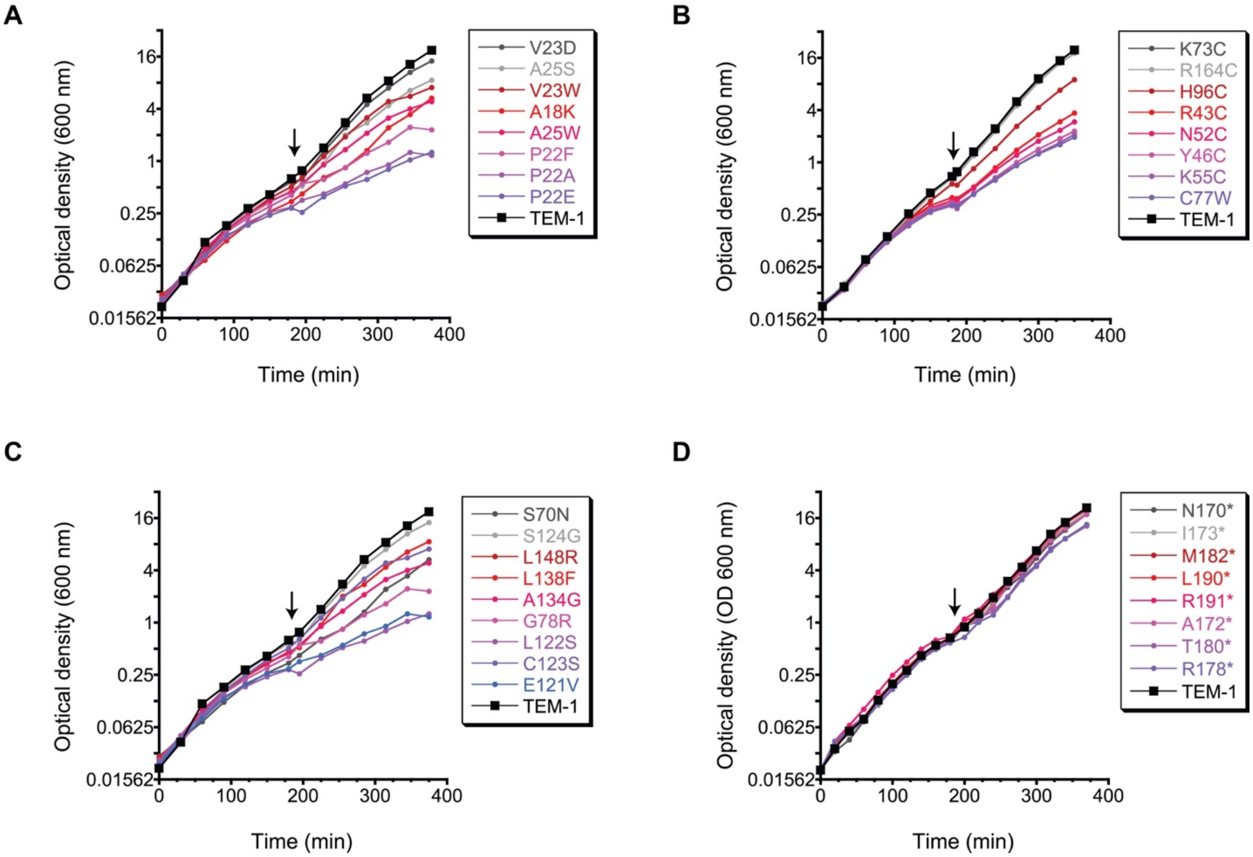


**Figure S8. Growth defects caused by select mutations.** Representative growth curves in monoculture growth assays (as measured by optical density at 600 nm) for cells expressing TEM-1 with (**A**) mutations in the signal sequence, (**B**) mutations involving cysteine, (**C**) mutations in $\alpha$-helices of the $\alpha$ domain, and (**D**) nonsense mutations. Growth assays performed as in **Fig S3B.** Arrows indicate the time of a 32-fold dilution into fresh media. OD measurements after this dilution are relative values in which the actual OD value is multiplied by the fold-dilution.

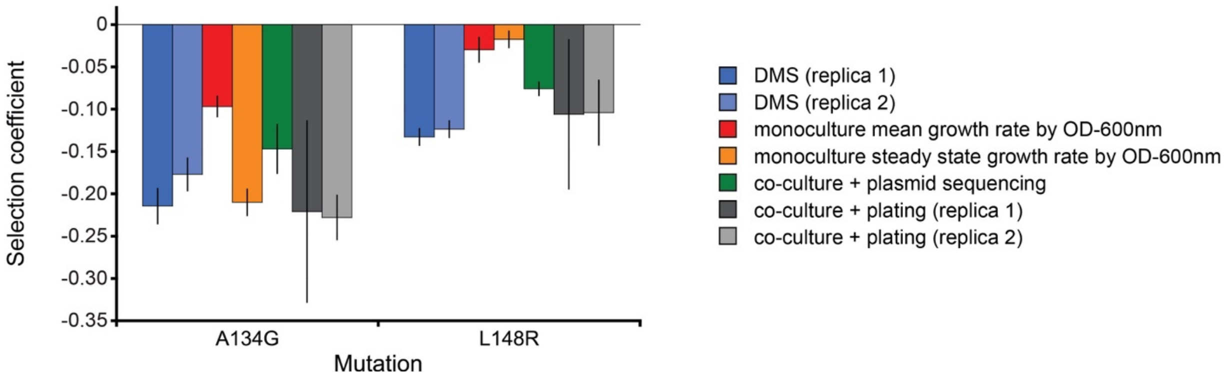


**Figure S9. Confirmation of the deleterious fitness effects of A134G and L148F by multiple methods.** The change in fitness caused by the mutations is indicated as measured by DMS (blue, two replica experiments, error bars represent 99% confidence intervals), mean growth rate of monoculture over 6 hours post-induction as monitored by OD-600nm (red, error bars represent 99% confidence intervals, *n*=3), steady state growth rate of monoculture between 4 and 6 hours post-induction as measured by OD-600nm (orange, error bars represent 99% confidence intervals, *n*=3), sequencing the plasmid isolated from co-culture at 6 hours post-induction and quantifying the sequencing chromatograms to determine the frequency of each allele (green, error bars represent 99% confidence intervals, *n*=3), and at 6 hours post-induction plating the co-culture on agar media lacking or containing ampicillin and determining the frequency of ampicillin resistant colonies (gray, two replica experiments, error bars represent standard error, *n*= 10 plates for each condition). The latter method works because A134G and L148R cause near total loss of TEM-1’s ability to provide Amp resistance.

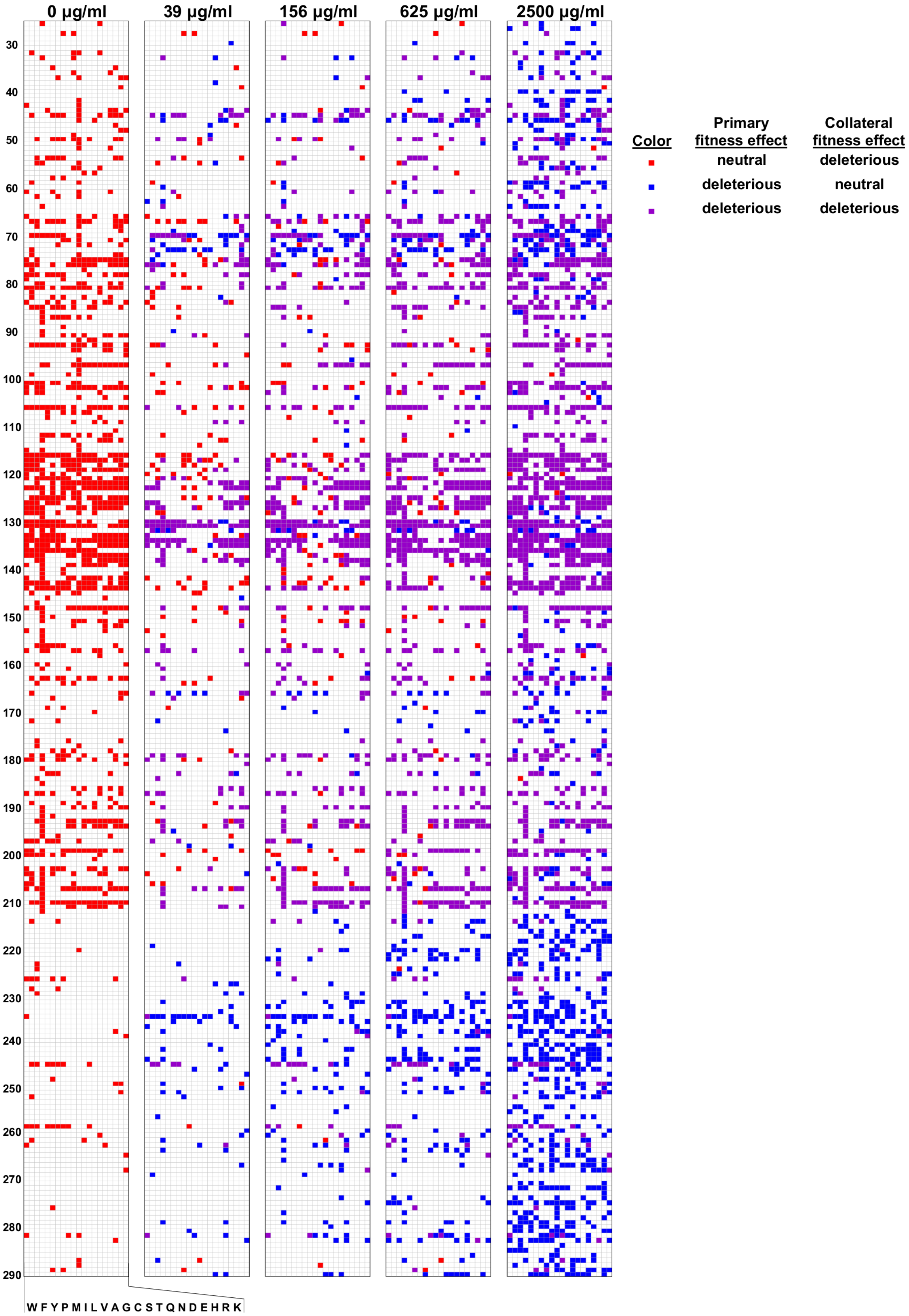
0 μg/ml
39 μg/ml
156 μg/ml
625 μg/ml
2500 μg/ml
30
40
50
60
70
80
90
100
110
120
130
140
150
160
170
180
190
200
210
220
230
240
250
260
270
280
290
W F Y P M I L V A G C S T Q N D E H R K
Color
Primary fitness effect
Collateral fitness effect
neutral
deleterious
deleterious
neutral
deleterious
deleterious

**Fig. S10. Sources of deleterious fitness effects.** The heat maps indicate whether a mutation had only collateral fitness effects (red), only primary fitness effects (blue), or both primary and collateral (purple). White indicates the data could not be assigned to any of these three categories, because of lack of data or because at least one fitness effect measurement could not be classified as deleterious or neutral (e.g. due to insufficient sequencing counts). The 99% confidence intervals were used along with the fitness values to assign mutations as deleterious, neutral, or neither (see Methods). Mutations assigned as deleterious had fitness values whose upper bounds were <0.98. Mutations assigned as neutral had fitness values whose lower bound were >0.98. Collateral fitness effects were measured in the absence of Amp in this study. Primary fitness effects were measured at 39, 156, 625, and 2500 µg/ml Amp from an experiment by Stiffler et al. (9).

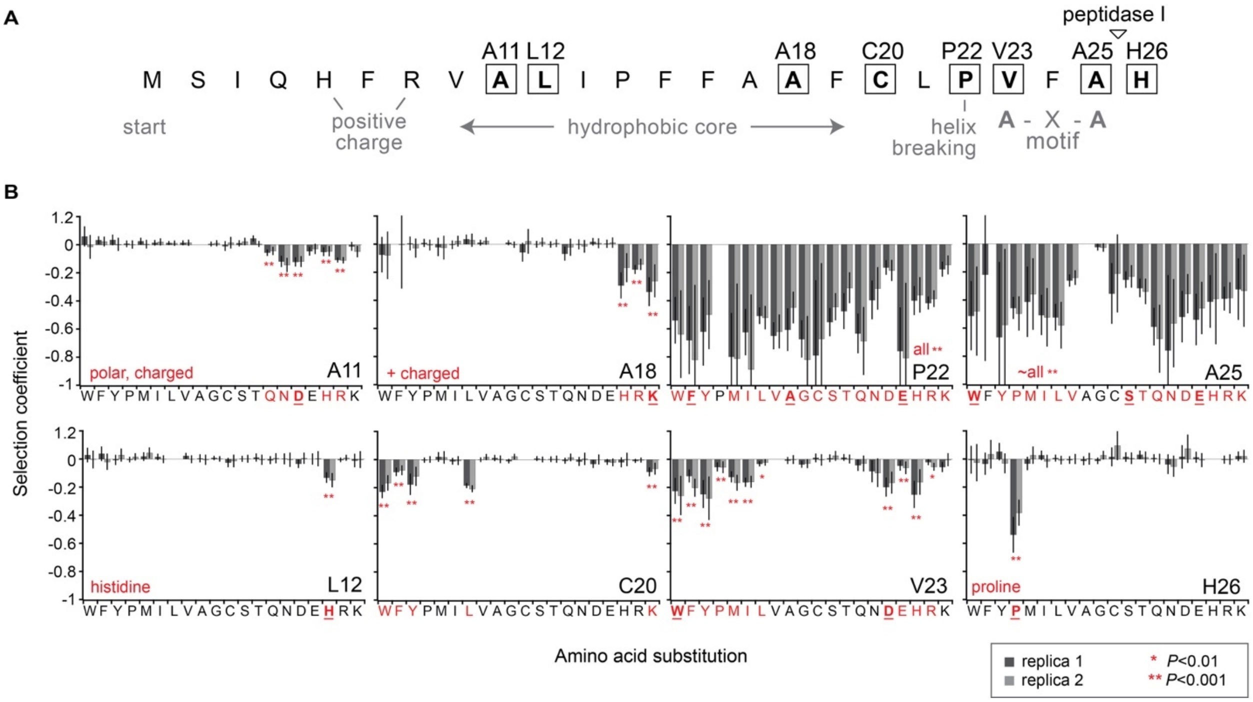


**Figure S11. Collateral fitness effects in the signal sequence.** (**A**) Key features of TEM-1's signal sequence. Residues with collateral fitness effects are boxed. The peptidase I cleavage site of the signal peptide is indicated by the triangle. (**B**) The effect of mutations at positions with collateral fitness effects. Mutations with effects in both replica experiments are indicated in red. Also indicated in red is the type of mutation causing the effect (if a pattern is apparent). The mutations selected for further study are bolded and underlined. *P*-value criteria are met for both DMS replica experiments.

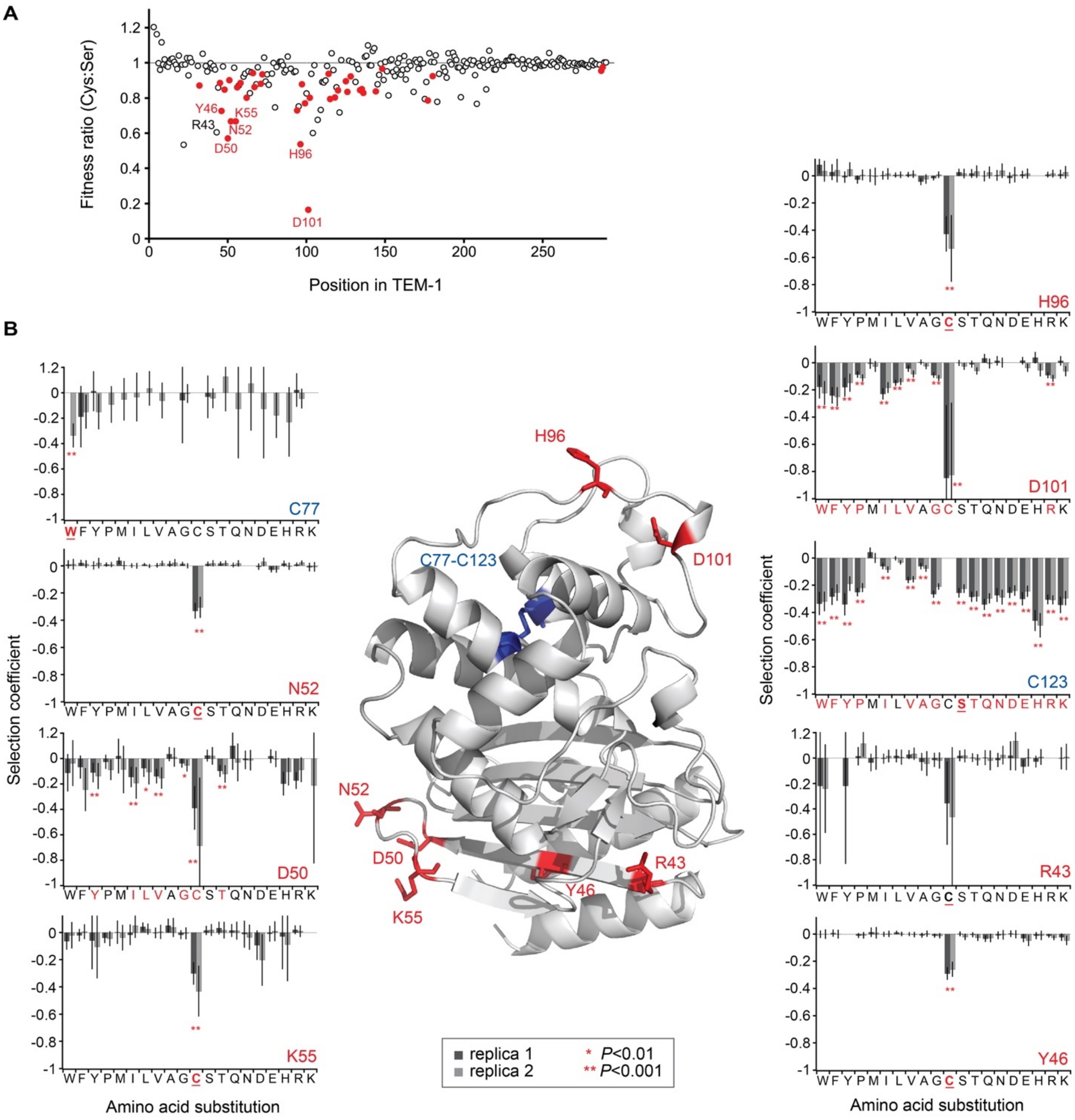


**Figure S12. Fitness effects of substitutions involving cysteine residues in TEM-1.** (**A**) Substitutions of Cys tend to be more deleterious than substitutions to Ser, but not after about position 210. Data in solid red correspond to positions where Cys mutations are deleterious ($P$<0.01) and Cys is more deleterious than Ser (assessed by propagation of the 99% confidence intervals as the error in the fitness values). (**B**) Fitness effects of mutations at cysteine residues (blue) and at positions where mutation to Cys is much more deleterious than mutation to serine (red). Mutations with fitness effects in both replica experiments are indicated in red. The mutations selected for further study are bolded and underlined. Data from two replica experiments are shown. Error bars are 99% confidence intervals. $P$-value criteria are met for both DMS replica experiments.

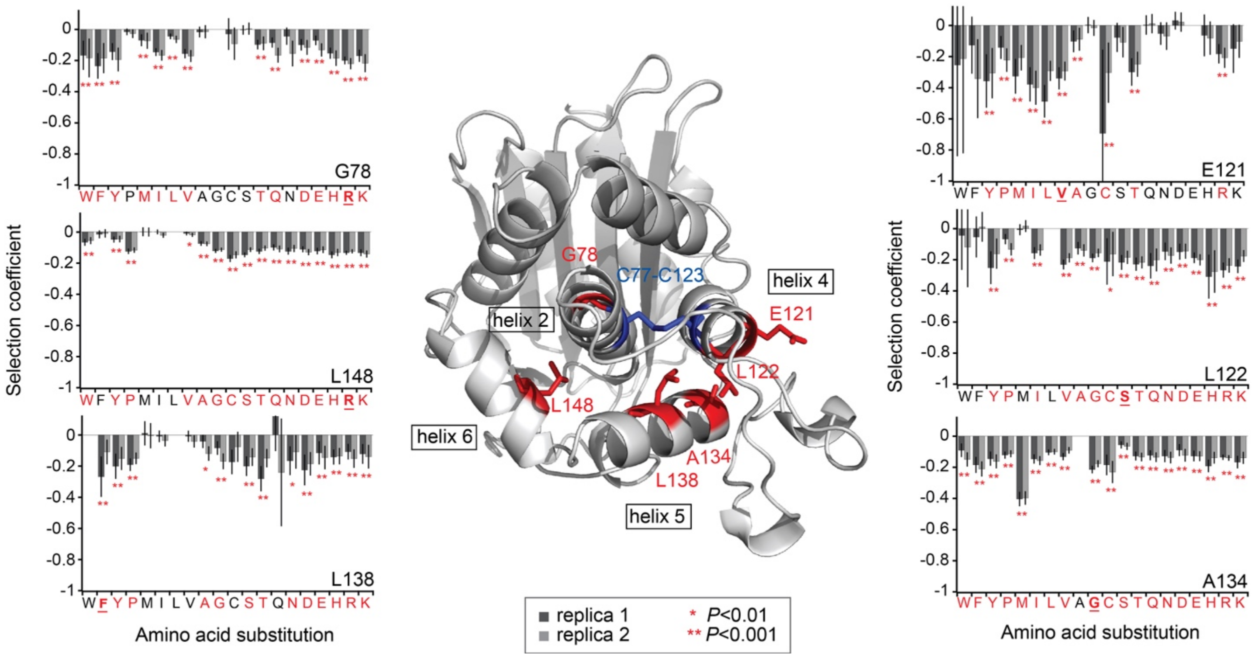


**Figure S13. Fitness effect of single-bp missense mutations in α-helices 2, 4, 5, and 6 in the α domain.** At the chosen sites, most mutations are deleterious. The α domain is in the foreground of the image. Mutations with fitness effects in both replica experiments are indicated in red. The mutations selected for further study are bolded and underlined. Data from two replica experiments are shown. Error bars are 99% confidence intervals. *P*-value criteria are met for both DMS replica experiments

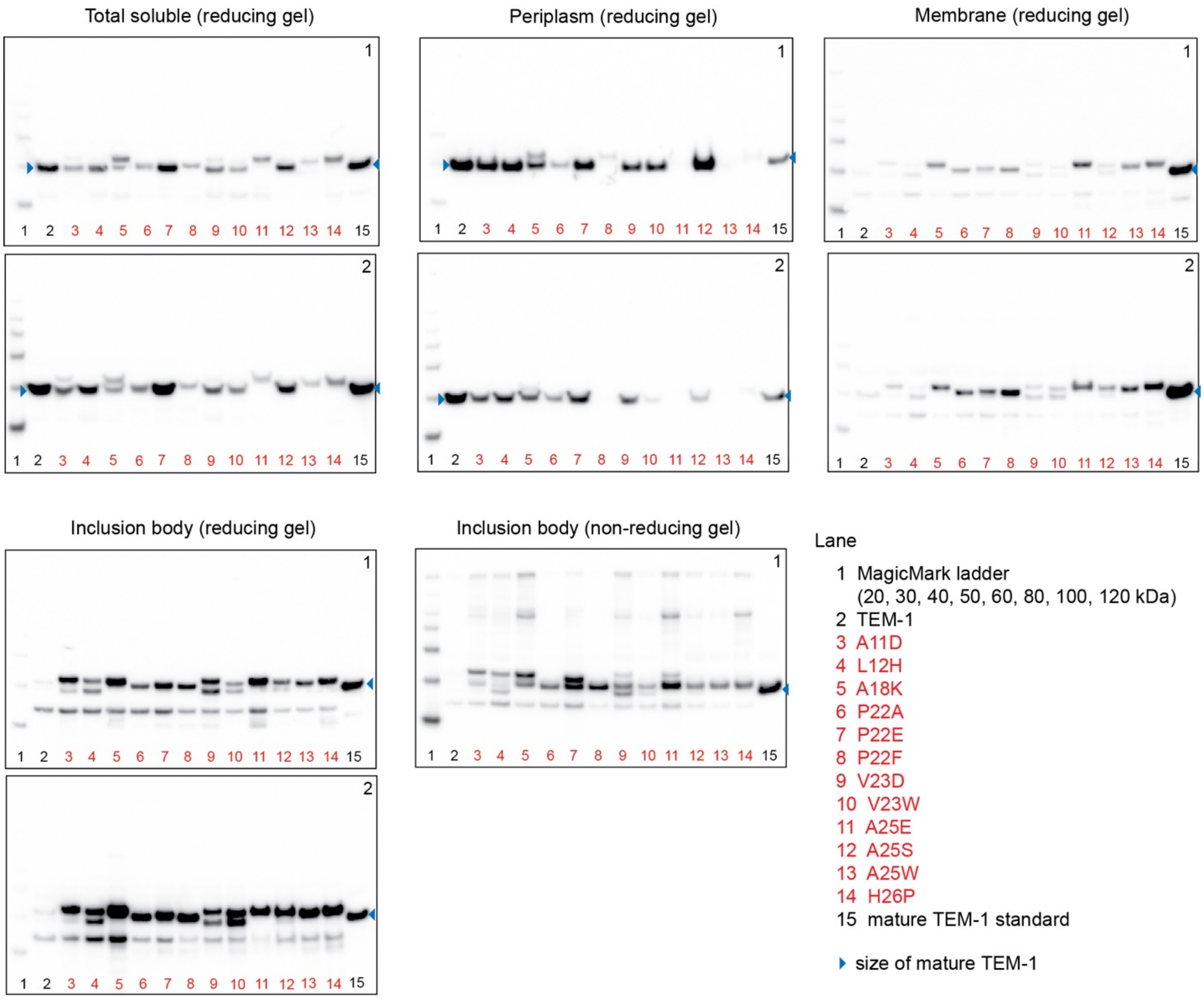


**Figure S14. Western blots of fractionated cells from replica cultures expressing TEM-1 with mutations in the signal sequence mutants.** The cell fraction and gel type are indicated at the top. The number in the upper right of each western is the replicate number of the cultures. Inclusion body and membrane samples are loaded at a higher level per OD unit than soluble samples in order to better see the mutations' effects. Deleterious mutations are labeled in red. Neutral mutations are labeled in black.

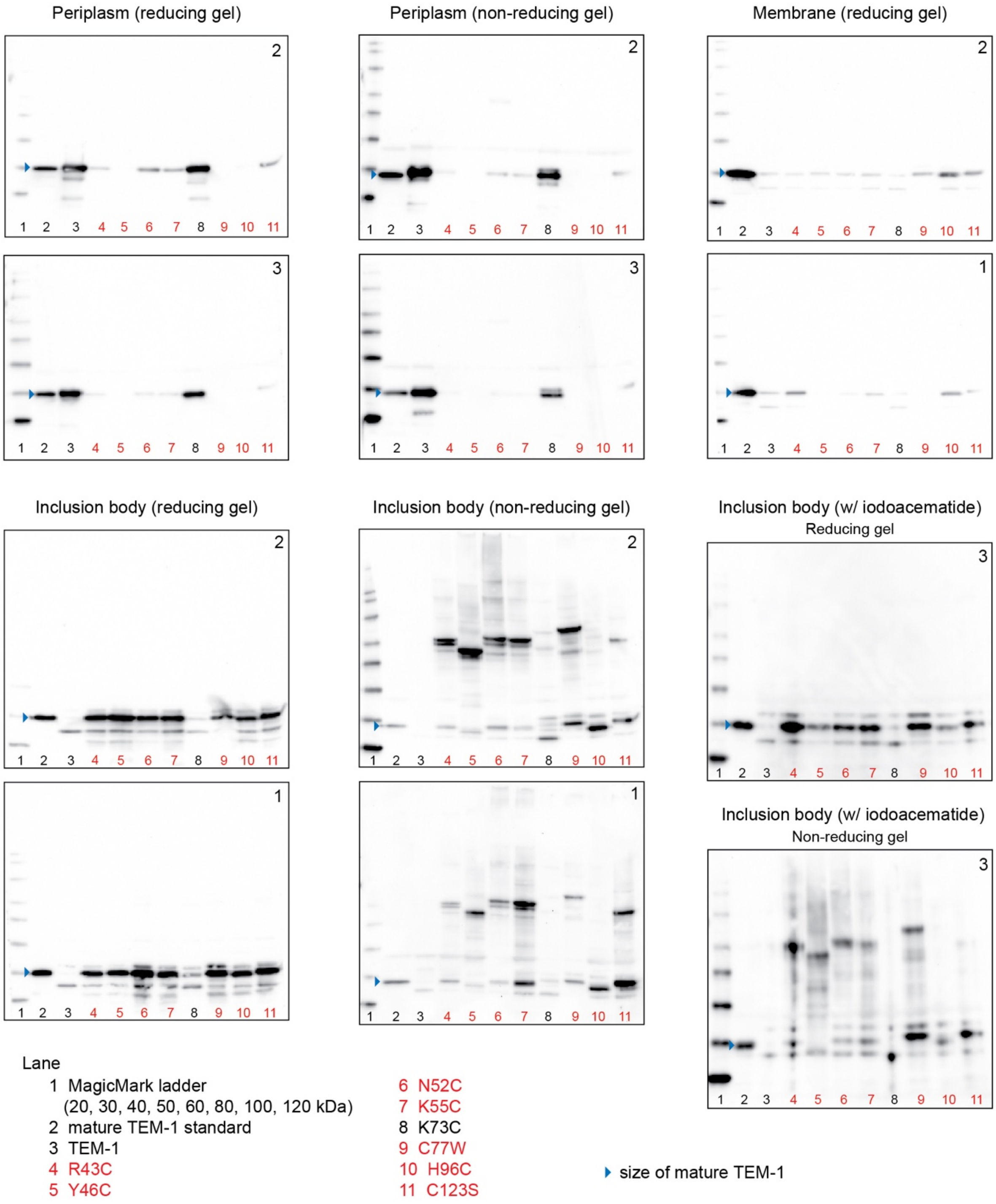


**Figure S15. Western blots of fractionated cells from replica cultures expressing TEM-1 with mutations involving cysteine residues.** The cell fraction and gel type are indicated at the top. The number in the upper right of each western is the replicate number of the cultures. Inclusion body and membrane samples are loaded at a higher level per OD unit than soluble samples in order to better see the mutations' effects. For periplasm samples, iodoacetamide was added to quench the cysteines as soon as the fraction was separated from the spheroplast fraction. For the inclusion body fraction with iodoacetamide, iodoacetamide was present in the cell lysis buffer. Deleterious mutations are labeled in red. Neutral mutations are labeled in black.

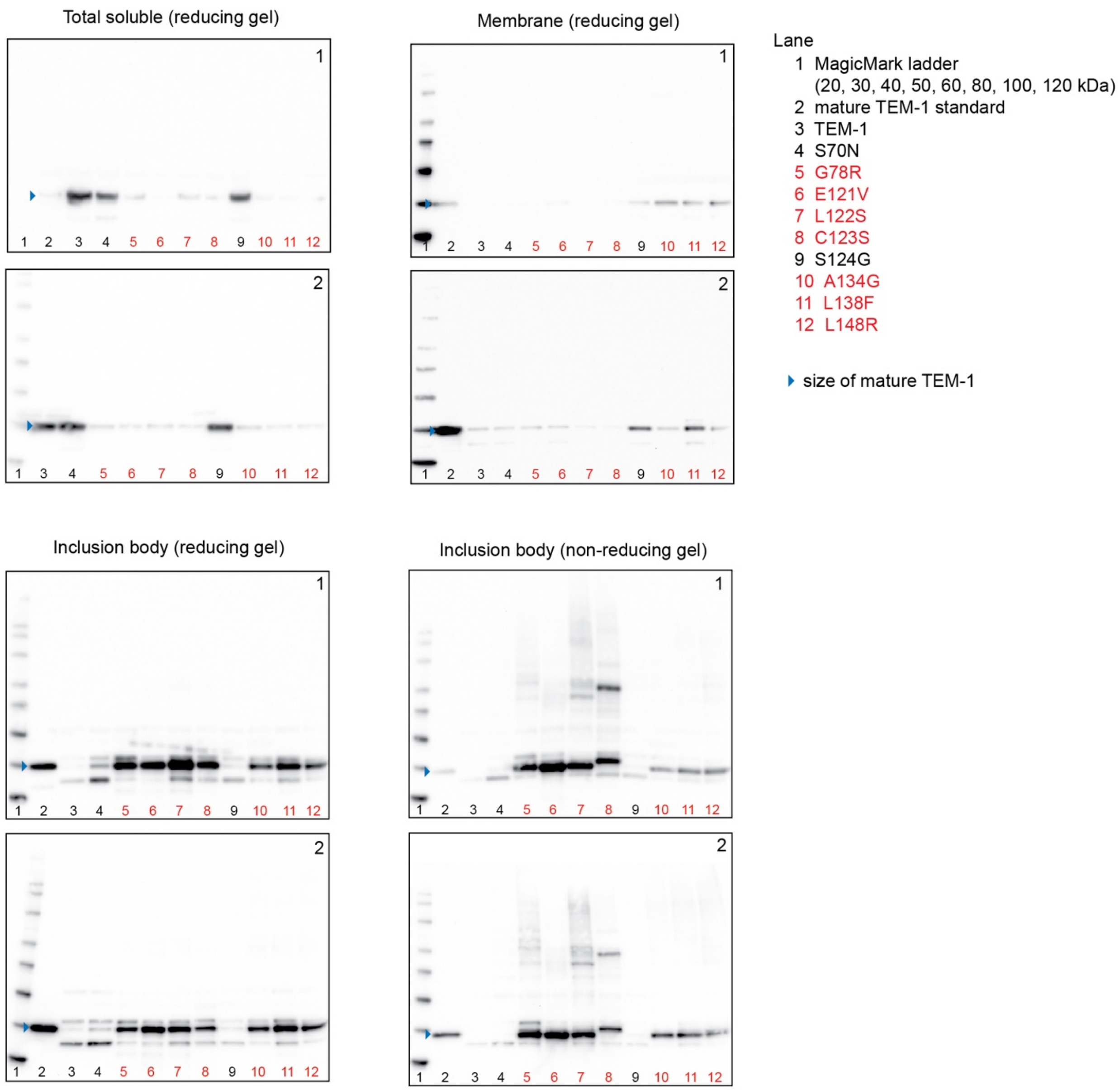


**Figure S16. Western blots of fractionated cells from replica cultures expressing TEM-1 with mutations in the α-helices of the α domain.** The cell fraction and gel type are indicated at the top. The number in the upper right of each western is the replicate number of the cultures. Inclusion body and membrane samples are loaded at a higher level per OD unit than soluble samples in order to better see the mutations' effects. Deleterious mutations are labeled in red. Neutral mutations are labeled in black.

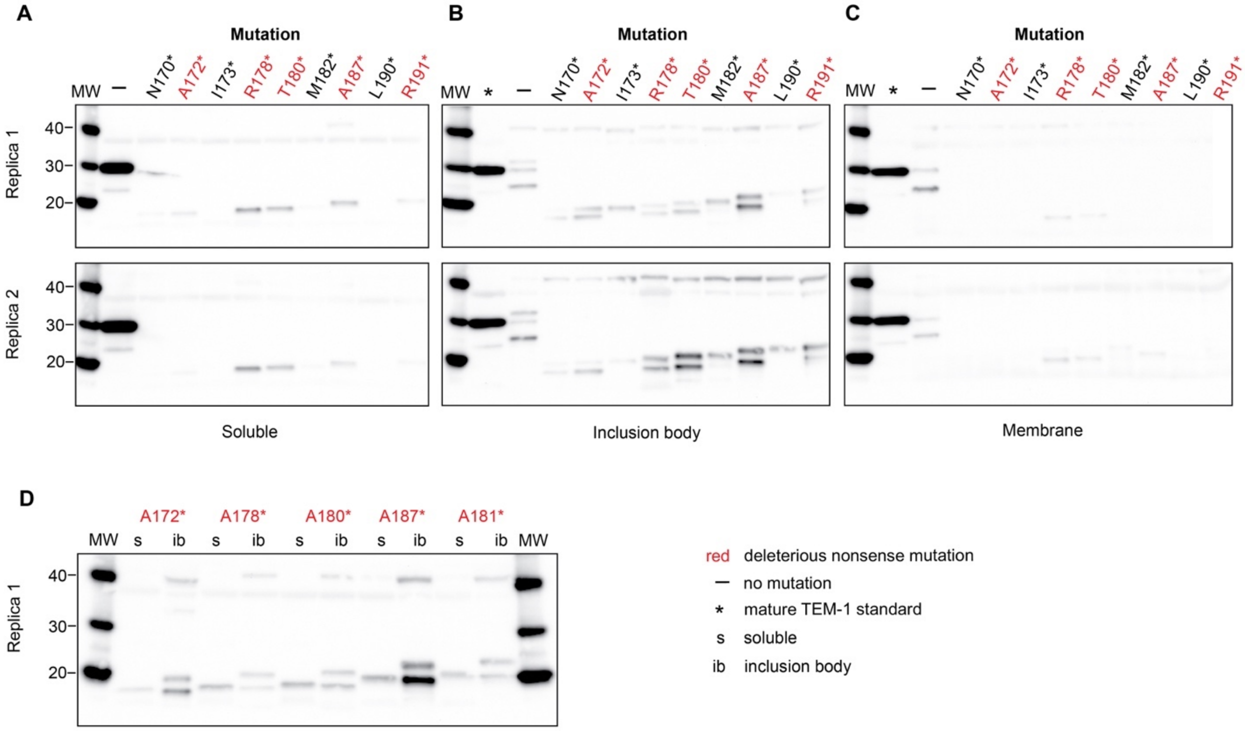


**Figure S17. Western blots of fractionated cells from replica cultures expressing TEM-1 with nonsense mutations.** (**A**) Total soluble fraction. (**B**) Inclusion body fraction. (**C**) Membrane fraction. Inclusion body and membrane samples are loaded at a higher level per OD unit than soluble samples in order to better see the mutations' effects. Deleterious mutations are labeled in red. Neutral mutations are labeled in black. (**D**) Western blot comparing the migration of TEM-1 fragments from the soluble and inclusion body fractions.

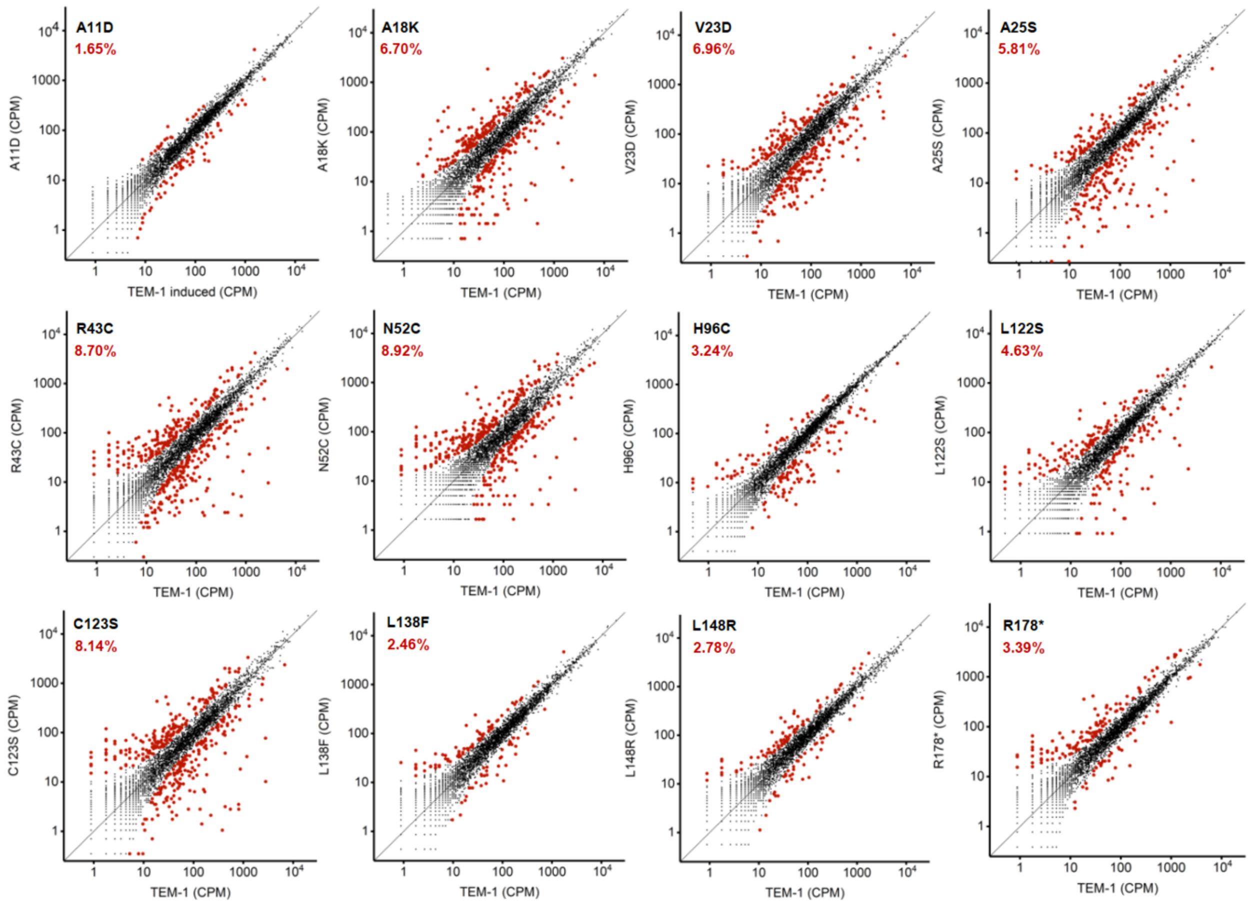


**Figure S18. Differential gene expression caused by mutations in TEM-1.** RNA-Seq results showing the mutation's effect on the transcriptome. Replica 1 data is shown for mutations with replica experiments. Data indicated with a larger, red circle indicates genes with at least a 2-fold change in expression and $P < 0.001$ ($Z$-test). The percentage of genes that meet these criteria is indicated on each plot. CPM, counts per million.

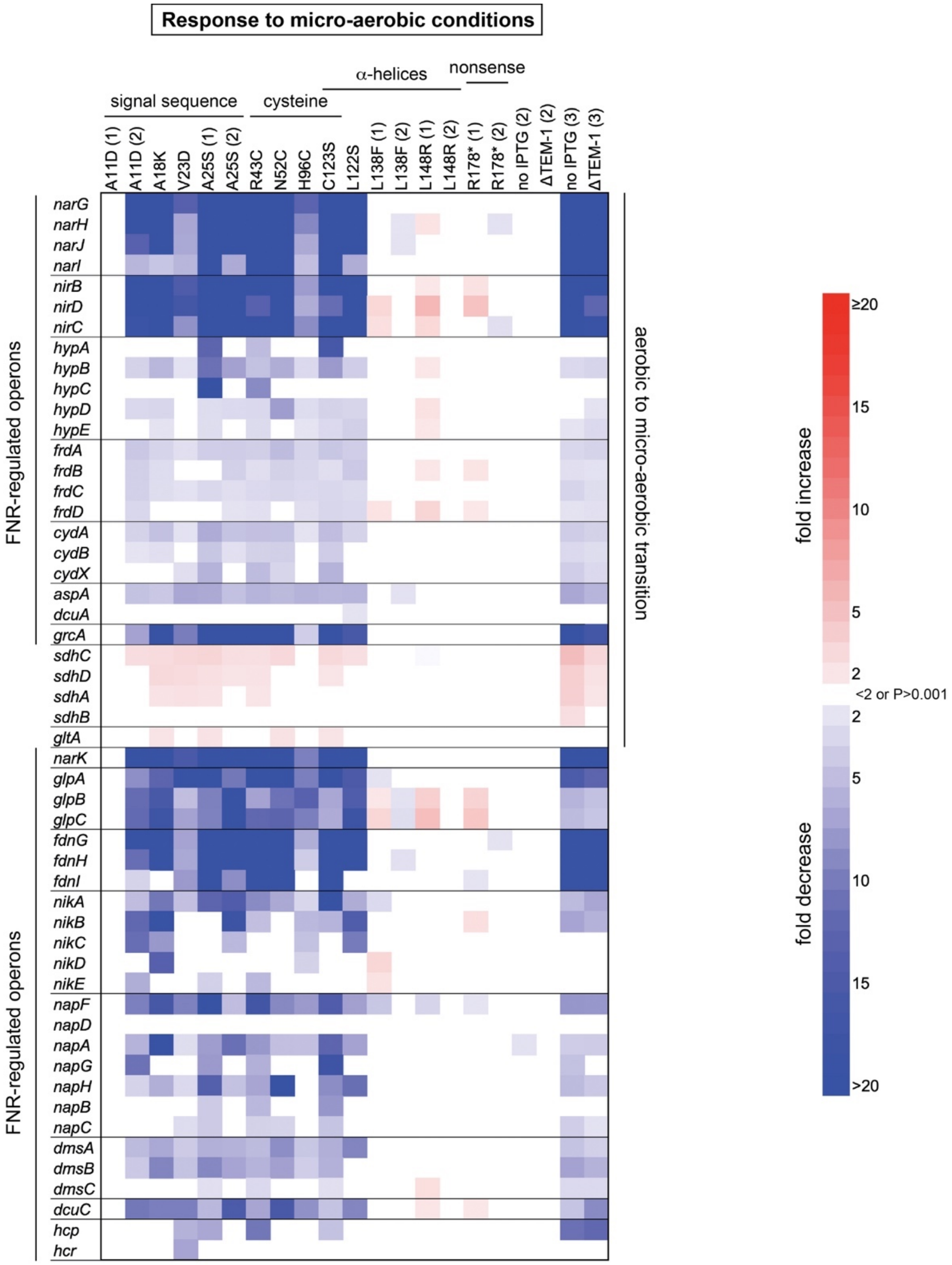


**Figure S19. Differential gene expression of select genes implicated in the transition from aerobic to micro-aerobic or anaerobic growth.** RNA-Seq experiments were performed on exponentially growing cells 5.5 hours post-induction of TEM-1. The heatmap shows changes in gene expression (relative to cells expressing TEM-1 lacking a mutation) that have a >2-fold change and *P* < 0.001. The number in parentheses after the mutation for samples with replica experiments indicate the day of the experiments. The control experiments in the last four columns correspond to cells lacking induction of TEM-1 expression (no IPTG) or cells lacking the TEM-1 gene but induced with IPTG (ΔTEM-1). Operons previously identified as induced or repressed when cells transitioned from aerobic to micro-aerobic conditions are listed in the top rows as indicated on the right (16). Previously-identified FNR-regulated operons are indicated on the left (15). FNR is required for the switch from aerobic to anaerobic growth.

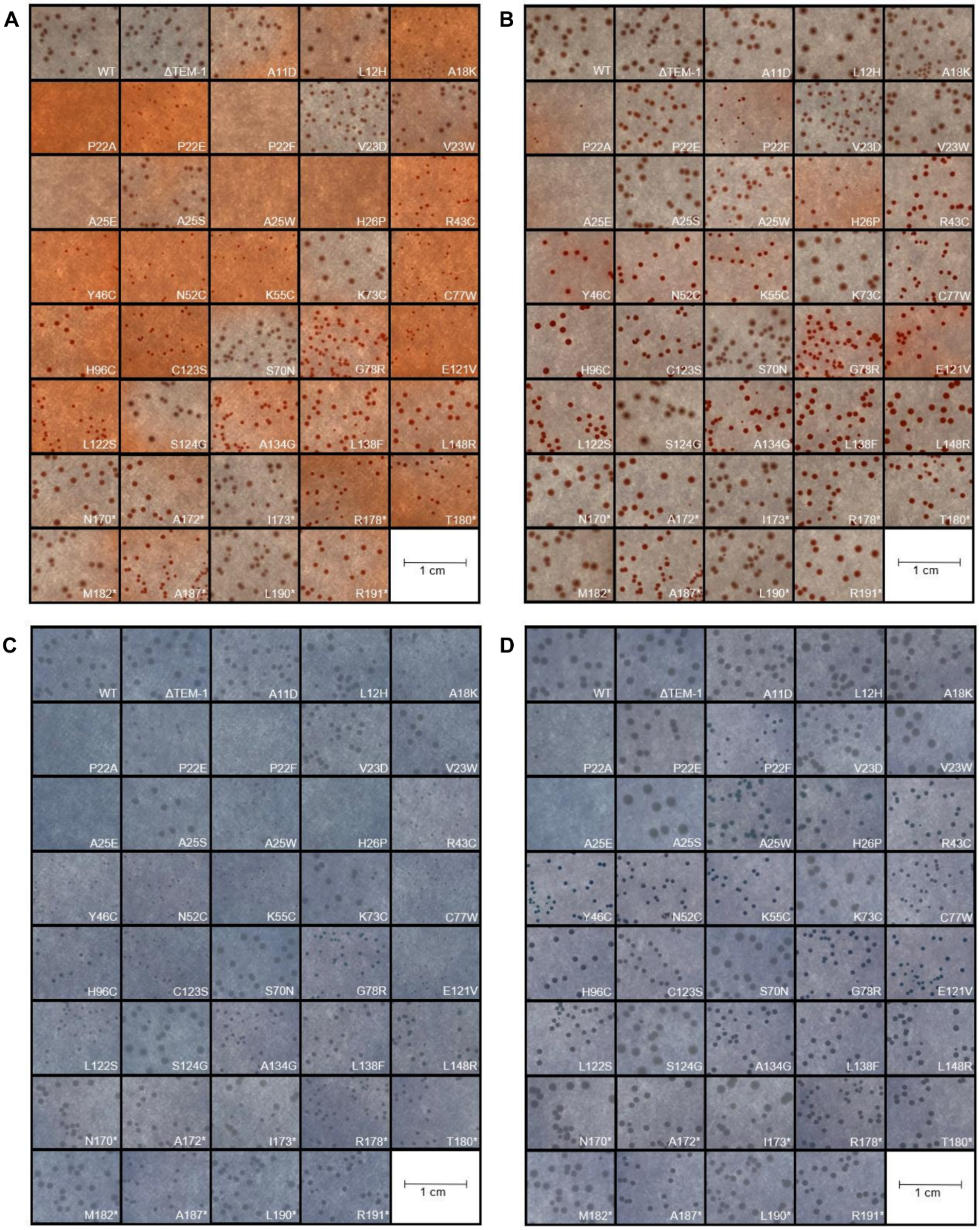


**Figure S20. Effect of mutations on the uptake of dyes diagnostic of EPS production.** Induced, exponentially growing cultures expressing TEM-1 and the indicated mutants were diluted and spread on LB-agar plates supplemented with 1 mM IPTG and 150 µg/ml Congo red (**A**,**B**) or 25 µg/ml toluidine blue-O (**C**,**D**). Plates were incubated for two days at 37°C and imaged at 1 day (A,C) and two days (B,D).

**Table S1. Fitness values for mutations studied individually**

| Mutation[a] | Fitness (-Amp)[b] | Fitness (+Amp)[c] | Fitness (39 μg/ml Amp)[d] | Fitness (156 μg/ml Amp)[d] | Fitness (625 μg/ml Amp)[d] | Fitness (2500 μg/ml Amp)[d] | SASA (%)[e] |
|---|---|---|---|---|---|---|---|
| **A11D** | 0.88 ± 0.01 | 0.18 ± 0.2 | - | - | - | - | |
| **L12H** | 0.86 ± 0.02 | 0.43 ± 0.04 | - | - | - | - | |
| **A18K** | 0.69 ± 0.10 | 0.35 ± 0.07 | - | - | - | - | |
| **P22A** | 0.50 ± 0.20 | 0.15 ± 0.02 | - | - | - | - | |
| **P22E** | 0.23 ± 0.06 | 1.05 ± 0.21 | - | - | - | - | |
| **P22F** | 0.30 ± 0.18 | 0.027 ± 0.005 | - | - | - | - | |
| **V23D** | 0.82 ± 0.04 | 0.25 ± 0.03 | - | - | - | - | |
| **V23W** | 0.77 ± 0.04 | 0.16 ± 0.02 | - | - | - | - | |
| **A25E** | 0.50 ± 0.10 | 0.15 ± 0.05 | - | - | - | - | |
| **A25S** | 0.75 ± 0.03 | 0.53 ± 0.10 | - | - | - | - | |
| **A25W** | 0.50 ± 0.04 | 0.032 ± 0.014 | - | - | - | - | |
| **H26P** | 0.56 ± 0.20 | 0.34 ± 0.05 | 0.93 ± 0.07 | 0.93 ± 0.02 | 0.70 ± 0.53 | 0.26 ± 0.16 | 45.93 |
| **R43C** | 0.62 ± 0.14 | 0.41 ± 0.03 | 1.04 ± 0.09 | 1.01 ± 0.08 | 0.67 ± 0.37 | -0.23 ± 0.02 | 30.86 |
| **Y46C** | 0.72 ± 0.04 | 0.030 ± 0.004 | 0.98 ± 0.03 | 0.12 ± 0.34 | -0.79 ± 0.19 | -1.04 ± 0.34 | 0 |
| **N52C** | 0.68 ± 0.03 | 0.55 ± 0.09 | 0.97 ± 0.05 | 0.99 ± 0.05 | 1.02 ± 0.03 | 0.19 ± 0.60 | 88.22 |
| **K55C** | 0.68 ± 0.17 | 0.51 ± 0.06 | 1.02 ± 0.02 | 0.96 ± 0.03 | 1.01 ± 0.05 | 0.05 ± 0.39 | 50.04 |
| **S70N** | 1.02 ± 0.02 | 0.0010 ± 0.0002 | -0.37 ± 0.04 | -0.29 ± 0.35 | -0.40 ± 0.28 | -0.29 ± 0.13 | 5.39 |
| **K73C** | 1.04 ± 0.11 | nd[f] | -0.20 ± 0.30 | -0.96 ± 0.59 | -0.67 ± 0.40 | -0.99 ± 0.54 | 0 |
| **C77W** | 0.66[g] | 0.0010 ± 0.0002 | -0.35 ± 0.28 | -0.41 ± 0.36 | -0.44 ± 0.24 | -0.58 ± 0.04 | 0 |
| **G78R** | 0.79 ± 0.03 | 0.0012 ± 0.0002 | -0.27 ± 0.37 | -0.73 ± 0.18 | -0.74 ± 0.09 | -0.97 ± 0.13 | 0 |
| **H96C** | 0.55 ± 0.14 | 0.56 ± 0.07 | 0.99 ± 0.07 | 0.99 ± 0.02 | 0.99 ± 0.02 | 0.18 ± 0.33 | 64.2 |
| **E121V** | 0.69 ± 0.06 | 0.007 ± 0.001 | 0.91 ± 0.06 | -0.26 ± 0.35 | -0.25 ± 0.69 | -0.05 ± 1.17 | 35.98 |
| **L122S** | 0.80 ± 0.03 | 0.034 ± 0.004 | 1.00 ± 0.07 | 0.86 ± 0.20 | -0.12 ± 0.27 | -0.50 ± 0.28 | 0 |
| **C123S** | 0.76 ± 0.04 | 0.14 ± 0.02 | 0.98 ± 0.03 | 0.96 ± 0.02 | 0.20 ± 0.46 | -0.33 ± 0.22 | 0 |
| **S124G** | 1.00 ± 0.01[h] | 1.00 ± 0.13 | 1.01 ± 0.08 | 1.00 ± 0.03 | 0.99 ± 0.03 | 1.01 ± 0.004 | 32.36 |
| **A134G** | 0.81 ± 0.05 | 0.16 ± 0.01 | 0.97 ± 0.02 | 1.02 ± 0.08 | 0.32 ± 0.90 | -0.20 ± 0.40 | 0 |
| **L138F** | 0.85 ± 0.20 | 0.0012 ± 0.0001 | 0.32 ± 0.54 | -0.57 ± 0.30 | -1.17 ± 0.18 | -0.77 ± 0.80 | 0 |
| **L148R** | 0.87 ± 0.01 | 0.0011 ± 0.0001 | -0.13 ± 0.22 | -0.91 ± 0.10 | -0.89 ± 0.55 | -1.10 ± 0.67 | 0.52 |
| **N170*** | 1.02 ± 0.02 | 0.0008 ± 0.0002 | -0.55 ± 0.02 | -0.50 ± 0.30 | -0.80 ± 0.09 | -1.18 ± 0.40 | 9.67 |
| **A172*** | 0.89 ± 0.05 | 0.0008 ± 0.0003 | 0.27 ± 0.24 | -0.70 ± 0.11 | -0.60 ± 0.56 | -0.73 ± 0.19 | 11.01 |
| **I173*** | 1.01 ± 0.03 | 0.0008 ± 0.0003 | 0.060 ± 0.59 | -0.12 ± 0.71 | -0.48 ± 0.26 | -0.36 ± 0.47 | 25.41 |
| **R178*** | 0.85 ± 0.02 | nd | 0.23 ± 0.32 | -0.80 ± 0.84 | -0.64 ± 0.85 | -0.68 ± 0.11 | 34.77 |
| **T180*** | 0.78 ± 0.07 | 0.0009 ± 0.0002 | -0.65 ± 0.01 | -0.75 ± 0.66 | -0.94 ± 0.18 | -0.83 ± 0.78 | 2.62 |
| **M182*** | 0.98 ± 0.03 | 0.0008 ± 0.0003 | 0.61 ± 0.55 | -0.40 ± 0.35 | -0.72 ± 0.58 | -0.56 ± 0.75 | 19.17 |
| **A187*** | 0.83 ± 0.04 | 0.0008 ± 0.0003 | -0.14 ± 0.71 | -0.97 ± 0.07 | -1.25 ± 0.42 | -1.30 ± 0.18 | 0 |
| **L190*** | 0.99 ± 0.03 | 0.0008 ± 0.0002 | 0.17 ± 0.18 | -0.60 ± 0.23 | -0.93 ± 0.14 | -0.60 ± 0.22 | 0 |
| **R191*** | 0.91 ± 0.01 | 0.0008 ± 0.0002 | 0.21 ± 0.35 | -0.42 ± 0.09 | -0.93 ± 0.65 | -0.46 ± 0.87 | 26.17 |

[a]Mutations are color-coded: black, signal sequence mutations; red, mutations involving cysteine; blue, mutations in α-helices of the α-domain; green, nonsense mutations; purple, mutation involving cysteine and in α-helices of the α-domain. The asterisk refers to a nonsense mutation.

[b]This study. Weighted mean of two replica DMS experiments with 99% confidence interval. Fitness is the mean growth rate of cells expressing the mutant protein in liquid media over ten generations post TEM-1 induction in the absence of Amp (relative to that of cells expressing TEM-1).

[c]Fitness is the relative ability of the gene to provide Amp resistance on agar plates as measured in Firnberg et al. (20) (with minor adjustments as explained in Steinberg and Ostermeier (21)). Error is the estimated uncertainty as explained in (20). Fitness values for nonsense mutations are specifically for the TAA codon.

[d]Experiment from Stiffler et al (9). Weighted mean of two replica experiments with 99% confidence intervals. Fitness is the mean growth rate of cells expressing the mutant protein in liquid media over 3-4 generations (relative to that of cells expressing TEM-1).

[e]Solvent-accessible surface area.

[f]Not determined

[g]Only one measurement.

[h]Excludes anomalous data of the GGC codon for Gly at this position (see **Fig. S3**)

**Captions for data files**

**Data S1. Excel file for DMS sequencing counts, fitness values, and associated statistics for collateral fitness effects.**

**Data S2. Excel file for DMS sequencing counts, fitness values, and associated statistics for primary fitness effects at 39 μg/ml ampicillin.**

**Data S3. Excel file for DMS sequencing counts, fitness values, and associated statistics for primary fitness effects at 156 μg/ml ampicillin.**

**Data S4. Excel file for DMS sequencing counts, fitness values, and associated statistics for primary fitness effects at 625 μg/ml ampicillin.**

**Data S5. Excel file for DMS sequencing counts, fitness values, and associated statistics for primary fitness effects at 2500 μg/ml ampicillin.**

**Data S6. Excel file for RNA-seq sequencing counts, fold-difference in expression, and associated statistics.**
